# The Dance of the Interneurons: How Inhibition Facilitates Fast Compressible and Reversible Learning in Hippocampus

**DOI:** 10.1101/318303

**Authors:** Wilten Nicola, Claudia Clopath

## Abstract

The hippocampus is capable of rapidly learning incoming information, even if that information is only observed once. Further, this information can be replayed in a compressed format in either forward or reversed modes during Sharp Wave Ripples (SPW-R). We leveraged state-of-the-art techniques in training recurrent spiking networks to demonstrate how primarily inhibitory networks of neurons in CA3 and CA1 can: 1) generate internal theta sequences or “time-cells” to bind externally elicited spikes in the presence of septal inhibition, 2) reversibly compress the learned representation in the form of a SPW-R when septal inhibition is removed, 3) generate and refine gamma-assemblies during SPW-R mediated compression, and 4) regulate the inter-ripple-interval timing between SPW-R’s in ripple clusters. From the fast time scale of neurons to the slow time scale of behaviors, inhibitory networks serve as the scaffolding for one-shot learning by replaying, reversing, refining, and regulating spike sequences.

## 1 Introduction

Immediately after watching a movie, we can often recall many scenes with great detail and even large stretches of dialogue. However, with the passing of time, our memories unfortunately may only preserve some aspects of the plot, along with a few memorable lines. This example illustrates how our memories operate on two different time scales. On the short time scale, we remember recent events clearly even with only a single presentation of a stimulus. On a longer time scale, we start to lose much of the finer details of any particular episode, yet we can still retain some aspects of it.

This observation, among others, led to the two-stage model of memory formation [Buzsáki, 1989, McClelland et al., 1995]. In the initial stage of memory formation, a labile or transient form of the memory is imprinted as some sort of engram onto hippocampal area CA3 rduring waking behavior. This initial acquisition is typically accompanied by a strong theta oscillation in the hippocampal local field potential (LFP). The second stage of memory formation occurs during consummatory or resting behaviors such as sleeping or eating. In this stage, strong correlated activity is initiated in CA3 which takes the form of a sharp wave (SPW) in the LFP. The SPW propagates through CA1 (as a sharp-wave ripple complex, SPW-R) and subsequently to the cortex for the long term consolidation of the memory. The spike train elicited by the stimulus during waking is replayed in compressed time during the SPW-R [Buzsáki, 2015].

The two-stage model of memory requires some mechanism for fast learning in a compressible format in the initial stage. In fact, the initial storage mechanism should cope with the fact that we often only receive a stimulus once. For example, behavioral experiments demonstrate our ability to recognize previously viewed visual stimuli such as pictures with high accuracy (> 90%) after only a single presentation [Standing et al., 1970]. Our accuracy in these tasks is much greater than chance levels with a steady decline with increasing time lag after presentation. Furthermore, this mechanism must allow for transfer from the temporary episodic form of memory into a consolidated semantic form, potentially through SPW-R compression. Thus, this hypothesized fast learning mechanism should allow for compressible spiking. Finally, this mechanism should be reversible. Spike sequences can be compressed in a reverse order immediately. For example, mice navigating a linear maze can reversibly compress spike sequences after only a single lap in a novel maze [Foster and Wilson, 2006].

What known feature of hippocampal functioning could endow us with our ability to learn immediately and satisfy all these constraints? Narrowing our focus to episodic memory formation yields one possible candidate: hippocampal internally generated theta sequences, or “time cells”. First discovered by [Pastalkova et al., 2008] in Hippocampal area CA1, these cells serve to partition time and are activated during episodic memory tasks [Pastalkova et al., 2008]. More recent findings demonstrate that time cells are also present in CA3 [Robinson et al., 2017], with temporally selective cells in the entorhinal cortex [Salz et al., 2016]. Like place cells, time cells also display phase precession [Pastalkova et al., 2008]. Time cells are critically dependent on the Medial Septum (MS) for normal functioning [Wang et al., 2015]. Here, we explore the idea that these time cells may serve as a temporal backbone to bind ongoing events to in a quick, online, compressible and reversible fashion.

Leveraging recent advances in the training of spiking neural networks [Nicola and Clopath, 2017, Sussillo and Abbott, 2009], we construct networks of phase precessing internally generated theta sequences (or time cells) based on interference theory [Burgess et al., 2007, O’keefe and Burgess, 2005, O’Keefe and Recce, 1993]. These time cells are generated through interference between an external GABAergic septal oscillation and intrinsic recurrent oscillation which operate at slightly different theta frequencies. Critically, both the external septal inputs and the recurrent theta oscillation are entirely mediated by inhibition and inhibitory connections. We refer to this network as the Septal-Hippocampal Oscillatory Theta (SHOT) network. The population activity for neurons in the SHOT network also displays theta frequency modulation that is directly inherited from the septal inhibition. Removal of the septal inputs can trigger compressed replay in the SHOT network. Further, we demonstrate that the time cells in the SHOT network can be used to quickly learn downstream sequences of spikes or behavioral quantities such as paths taken during navigation. When learning spike trains, the resulting learning rule is a local Hebbian rule and allows for single-trial learning. Surprisingly, compressed replay in conjunction with the Hebbian learning rule segregates spiking into compressed gamma-assemblies while application of septal inhibition after learning yields gamma assemblies nested on a theta oscillation. Reverse compressed replay, as seen during quiet awake periods, can also be elicited by using a second population of interneurons. Finally, we demonstrate that this mechanism for triggering compression and rapid learning is compatible with the synthesis of sharp-wave ripples. Using the SHOT network as a model for CA3, and a downstream network as a model for CA1, we demonstrate how CA1 inhibitory neurons refine gamma assembly width and prevent assemblies from interfering with each other. Further, we find that allowing the CA3 inhibitory neurons to influence the probability of SPW-R initiation prevents the fragmentation of memories and results in the formation of ripple clusters. Our results demonstrate concrete mechanisms for how inhibitory networks can gate, control, transform, and refine spike sequences that occur during both sharpwave ripples and the theta oscillation to promote rapid learning.

## 2 Results

### 2.1 Generating Internally Generated Theta Assemblies with Network Based Oscillatory Interference

In order to construct a plausible sequence of time cells, we use the experimental literature to constrain and inform our initial network. First, time cells critically depend on inputs from MS as injections of the GABA agonist muscimol into MS disable time-field expression [Wang et al., 2015]. The septal inputs consist of GABAergic, cholinergic, and glutamatergic connections. The GABAergic connections innervate inhibitory interneurons exclusively [Freund and Antal, 1988, Gulyás et al., 1990, Dragoi et al., 1999] in their downstream hippocampal targets while the cholinergic inputs are more widespread and non-selective [Frotscher and Léránth, 1985, Freund and Antal, 1988, Lewis and Shute, 1966]. Furthermore, many of these neurons fire bursts of action potentials at a theta frequency locked in phase with the hippocampal theta oscillation [Joshi et al., 2017]. These experimental results yield the first component of the Septal-Hippocampal Oscillatory Theta (SHOT) network, which is an inhibitory oscillation from the MS:

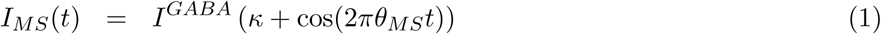

The parameter *θ_MS_* is the theta frequency oscillation of rhythmic GABAergic interneurons in the MS, *κ* ≥ 1 is a parameter that controls the baseline inhibition levels arriving from the MS, and *I^GABA^* controls the amplitude of the oscillation.

Next, time cells generate a sequence of firing fields that operate on a behavioral time scale, while the individual neurons oscillate at a theta frequency inside the firing field [Pastalkova et al., 2008]. Their rate of oscillation is slightly faster than the theta oscillation frequency of the local field potential and as a result they precess in phase [Pastalkova et al., 2008]. One potential mechanism underlying this phenomenon is the emergence of time cells through the interference pattern of two GABAergic septal neurons firing at slightly different theta frequencies [Hasselmo and Stern, 2014]. Alternatively, phase precessing firing fields may be generated by local circuitry involving pairs of neurons within the hippocampus receiving septal inhibition [Chadwick et al., 2016]. Unlike previous work, we will take the perspective that the firing fields are a network phenomenon created by the interference between septal and intra-hippocampal inhibitory oscillators.

Finally, inspired by experimental and theoretical findings that demonstrate the rhythm generation capabilities of the hippocampus [Buzsáki, 2002, Buzsáki et al., 1986, Kramis et al., 1975, Konopacki et al., 1987, Goutagny et al., 2009, Ferguson et al., 2017, Amilhon et al., 2015], we hypothesized that septal inputs could interact with a recurrently coupled population of interneurons that was oscillating at its own internal frequency, *θ_INT_* (INT for Interneuron), to generate a robust interference pattern. To avoid exhaustive parameter searches in determining how a network of neurons should be coupled, we leveraged recent theoretical advances in training recurrent spiking neural networks ([Nicola and Clopath, 2017, Sussillo and Abbott, 2009]). We successfully FORCE trained a population of neurons to oscillate at its own internal frequency, *θ_INT_*, while simultaneously receiving septal inhibition of frequency *θ_MS_* (Figure 1A, Supplementary Figure S1). Our network consists of *N_E_* = 2000 excitatory and *N_I_* = 2000 inhibitory leaky-integrate-and-fire (LIF) neurons. The training is constrained to only inhibitory connections (see Supplementary Figure S1, Materials and Methods Section: FORCE Training a Balanced E/I Network with Dales Law). The excitatory neurons receive inputs from the inhibitory neurons, but do not project any excitatory connections. They serve as readouts for the inhibitory neurons in the SHOT network and receive constant background currents to trigger spiking. Finally, the excitatory neurons in the SHOT network do not receive septal inhibition.

**Figure 1:**
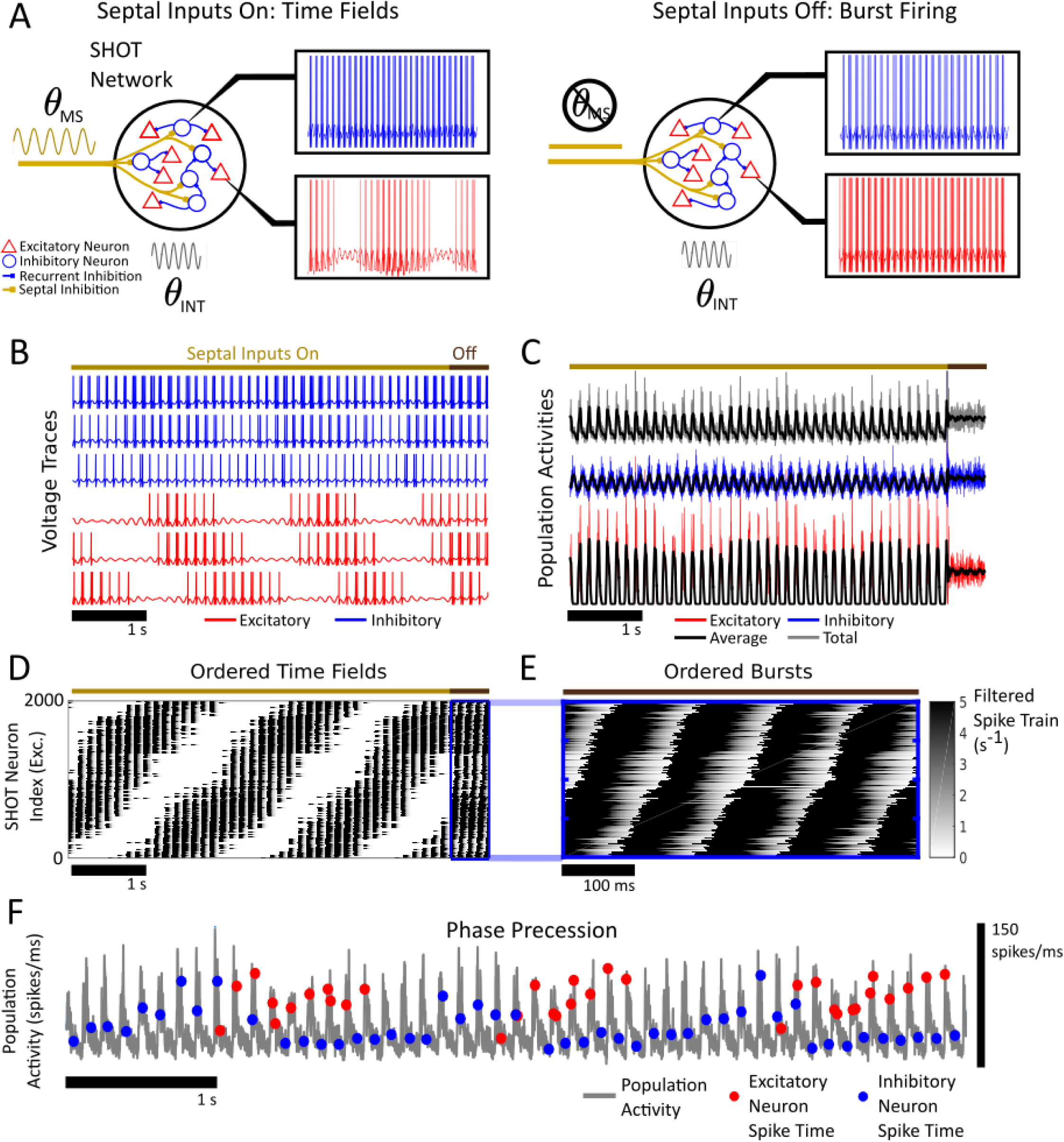
FORCE Training the SHOT network. **(A)** A schematic for the behavior of the Septal-Hippocampal Oscillatory Theta (SHOT) network. (Left) When GABAergic inputs (gold) from the medial septum (MS) are present, firing fields emerge in the excitatory SHOT neurons on the behavioral time scale (red) while the inhibitory SHOT neurons (blue) fire theta-frequency bursts. (Right) When septal inputs are absent, all SHOT neurons fire bursts. The inhibitory SHOT neurons are trained to oscillate at a *θ_INT_* frequency. No excitatory connections in the SHOT network are present. The excitatory neurons receive a constant background current to elicit firing. **(B)** The voltage traces for 3 excitatory neurons (red), 3 inhibitory neurons (blue) for the FORCE trained SHOT network for 5.5 seconds of simulation time. The overline denotes the presence (gold, first 5 s) or absence (brown, last 500 ms) of septal inhibition. The network is trained to produce its own internal oscillation at a frequency of *θ_INT_* = 8.5 Hz simultaneous to the application of oscillatory septal inhibition with a frequency of *θ_MS_* = 8 Hz. The result is an interference pattern generated due to the septal and the recurrent inhibition generated by the network. **(C)** The population activities of spike times of the SHOT network are computed as histograms by binning over 1 ms bins for the excitatory (blue), inhibitory (red) and total (grey) populations in addition to the smoothed time averages (black). The smoothing was performed with a 20 ms box filter. The network firing statistics display a prominent theta oscillation in the presence of septal inhibition but not in their absence. Furthermore, the dynamics of the population activities display negligible characteristics of an interference pattern and primarily follow the external septal inputs. **(D)** The excitatory SHOT neurons are sequentially active. The firing fields are plotted as the synaptically filtered spike trains ***r***(*t*) with the synaptic filters *τ_R_* = 2 ms and *τ_D_* = 20 ms. Darker colours indicate a larger value of ***r***(*t*). For visual clarity, we have thresholded the upper value of ***r***(*t*) at 5 Hz. Removal of the septal inputs results in the recurrent dynamics taking over and the firing of bursts in the last 500 ms seconds of the simulation. **(E)** A zoomed in segment consisting of 500 ms without septal inputs present. The order of bursts in the recurrent dynamics determines the ordering of time fields and vice-versa. To trigger bursts, the excitatory neurons receive an extra current (valued at 20 pA) when the septal currents are off. **(F)** Phase precessing spikes for an excitatory neuron (red points, spike times) and an inhibitory neuron (blue points, spike times) in the network in reference to total population activity (in grey, both excitatory and inhibitory SHOT neurons) with septal inputs present. The population activity is measured as in (C), a histogram with 1 ms time bins for 6 seconds with septal inhibition present.

After training, we examined the activity of neurons in the SHOT network. When septal inhibition is present, the inhibitory SHOT neurons fire bursts periodically for every theta cycle with some heterogeneity from neuron to neuron (Figure 1B). The excitatory neurons however fire isolated firing fields on a behavioral time scale (Figure 1B). Furthermore, population activities of both excitatory and inhibitory neurons displayed very minimal interference patterns (Figure 1C) and were tightly locked to the septal inhibition frequency. Any interference pattern in the population activities present will decrease with larger network sizes (see Materials and Methods: Interference Based Control of the Population Activity by Septal Inputs). The phase locking of population activities to septal inhibition is a consequence of interference theory (see Materials and Methods: Interference Based Control of the Population Activity by Septal Inputs). Thus, in the normal operating regime under which the network was trained, we observed the emergence of stable, ordered firing fields (Figure 1D). Finally, by using either the MS input oscillation (not shown) or the population activity (Figure 1F) as a reference theta oscillation, both excitatory and inhibitory neurons precess in phase due to the interference between the inhibitory and septal oscillations. This is a standard result from interference theory [Burgess et al., 2007, O’keefe and Burgess, 2005, O’Keefe and Recce, 1993].

### 2.2 Removing Septal Inputs Can Trigger Compressed Replay

The SHOT network was trained to oscillate at a *θ_INT_* frequency while simultaneously receiving septal inhibition at a frequency of *θ_MS_*. Thus, we investigated how the network would extrapolate to removal of septal inhibition (Figure 1A, right). We found that removing the septal inputs transformed the time fields into sub-threshold oscillations in the excitatory neurons (Supplementary Figure S2A). In the absence of septal inputs, each excitatory neuron receives increased inhibition from the interneurons in the SHOT network and does not spike. However, the phase of subthreshold oscillation in the excitatory SHOT neurons mirrors their time-field ordering when septal inhibition was present. This sub-threshold compression is a consequence of interference theory (see Materials and Methods: Interference Based Compression of Spike Sequences). The increased inhibition in the excitatory neurons is due to the inhibitory neurons in the SHOT network no longer receiving septal inhibition. Thus, in the absence of septal inputs the excitatory population is normally not spiking. However, if the excitatory neurons are provided with an additional excitatory current when septal inhibition is removed, bursts are triggered (Figure 1B,D-E, Supplementary Figure S2B,. The extra current provided to the excitatory neurons can be thought of as a proxy for recurrent excitation within the hippocampus. The excitatory SHOT neurons now fire ordered bursts on a compressed time scale in the absence of septal inhibition. These bursts are in the same order on as the firing fields were in the presence of septal inhibition. This demonstrates a concrete mechanism by which inhibitory networks orchestrate the timing of compressed replay. The genesis of spiking, on the other hand, is regulated by other mechanisms such as recurrent excitation. Additionally, the measured population activities show minimal signs of *θ_INT_* in the absence of septal inhibition. Rather, the population activities fluctuated around a constant value (Figure 1C). The lack of *θ_INT_* in the population activity is also a consequence of interference theory (see Materials and Methods: Interference Based Control of the Population Activity by Septal Inputs). Thus, in the presence of septal inhibition, ordered firing fields emerge along with theta modulation (*θ_MS_*) of the population activity. In the absence of septal inhibition, the time-field is compressed down to a burst with an oscillation frequency of *θ_INT_* while the population activity does not display *θ_INT_* rhythmicity.

As the extra current served as a proxy for recurrent excitation, we considered directly coupling the excitatory SHOT neurons with dense recurrent excitation. Activation of a small subset of these excitatory neurons is sufficient to trigger compressed, ordered firing (Supplementary Figure S2D). However, activation of too many neurons can lead to pathological, high frequency firing due to run-away excitation (not shown). Thus, for simplicity and stability, we apply extra current instead of recurrent excitation in all subsequent simulations.

Finally, in place of recurrent excitation, disinhibition of the excitatory neurons in the SHOT network can also trigger ordered compressed bursts (Supplementary Figure S2C). In particular, transiently increasing septal inhibition instead of removing it can trigger compressed bursts and compressed replay of the time-fields (Supplementary Figure S2C). Our model offers theoretical validation of a disinhibitory mechanism for SPW-R generation involving the silencing of axo-axonic (AAC) cells [Viney et al., 2013, Somogyi et al., 2014]. Indeed, in [Viney et al., 2013], the authors find a subset of septal GABAergic interneurons that project to AAC cells in CA3 that increase their population activity during SPW-R. This causes an inhibition of AAC cells which preferentially inhibit the axon-initial segment of pyramidal neurons. The inhibition of these AAC cells was hypothesized to be a necessary condition for SPW-R generation [Viney et al., 2013].

Thus, removing septal inputs can trigger burst firing where the order of bursts in the network mirrors the order of firing-fields when septal inputs were present, only on a compressed time scale. The bursts can be triggered by either recurrent excitation and removal of septal inhibition, or disinhibition by applying extra septal inhibition to the SHOT interneurons. The order of bursts is regulated exclusively by the interneurons in the SHOT network while the induction of bursting is regulated by either recurrent excitation or disinhibition.

### 2.3 Firing Field Generation is Robust

If the time cells form a backbone for compressible learning, the firing field formation should be robust. Thus, we investigated how the SHOT network would respond to non-ideal conditions. For example, our network was trained under the septal input frequency of *θ_MS_* = 8 Hz with the constant background currents to neurons set at a specific level. It is unclear how the network would extrapolate to other frequencies or background levels of excitation. If the SHOT network was operating as a perfect linear oscillator, we should expect that the time-field periodicity scale like |*θ_MS_* − *θ_INT_*|^−1^ near *θ_INT_*, while the population activity to have a periodicity of 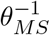. Furthermore, interference theory predicts that the time-field order reverses when *θ_MS_* > *θ_INT_* (See Materials and Methods: Interference Based Compression of Spike Sequences). Finally, we should also expect time-fields to also emerge near the harmonics of *θ_INT_*. To test this hypothesis, we slowly swept the frequency of the septal inputs from 2 Hz to 12 Hz and monitored the time-field periodicity, sequential order, and the population activity of the SHOT network (Supplementary Figure S3). We found that the SHOT network behaves as a nearly perfect linear oscillator for all input frequencies tested. The firing fields also reverse near the harmonics of *θ_INT_* (Supplementary Figure S3A). For all frequencies tested, the septal frequency directly determines the frequency of the population activity, as predicted from interference theory. Finally, we also considered amplitude modulations to both the background current to the inhibitory SHOT neurons, and the inhibitory septal currents. Ramping the excitatory current to the inhibitory population has no effect on the population activity, and a small yet statistically significant effect on the periodicity of the interference pattern (Supplementary Figure S4). Ramping the septal amplitude (*I^GABA^*) however has a large and statistically significant effect on the periodicity of the interference pattern and a small yet significant effect on the frequency of the population activity (Supplementary Figure S4). Thus, the network is sensitive to modulations of the septal inhibition amplitude. The interference pattern breaks down at about 1.5 times the baseline the baseline amplitude.

### 2.4 Reverse Compressed Replay

Given that the internally generated theta sequences were compressible and robust, we investigated if a reverse compression mechanism also existed within the framework of interference theory, as opposed to other mechanisms for reverse replay [Roach et al., 2018, Buzsáki, 2015]. We discovered three potential mechanisms for reverse compression (Figure 2). First, if the septal inputs are higher in frequency than the recurrent frequency of the SHOT network (*θ_MS_* > *θ_INT_*), than the order of firing fields reverses (Figure 2A-E). We refer to this mechanism as Mirror Reversion as removal of the septal inputs reverses the order of bursts during compressed replay (Figure 2C). However, this is not a plausible mechanism as the septal frequency requirement also results in phase regression (Figure 2E) and reversed theta sequences (Figure 2D) as opposed to phase precession.

**Figure 2:**
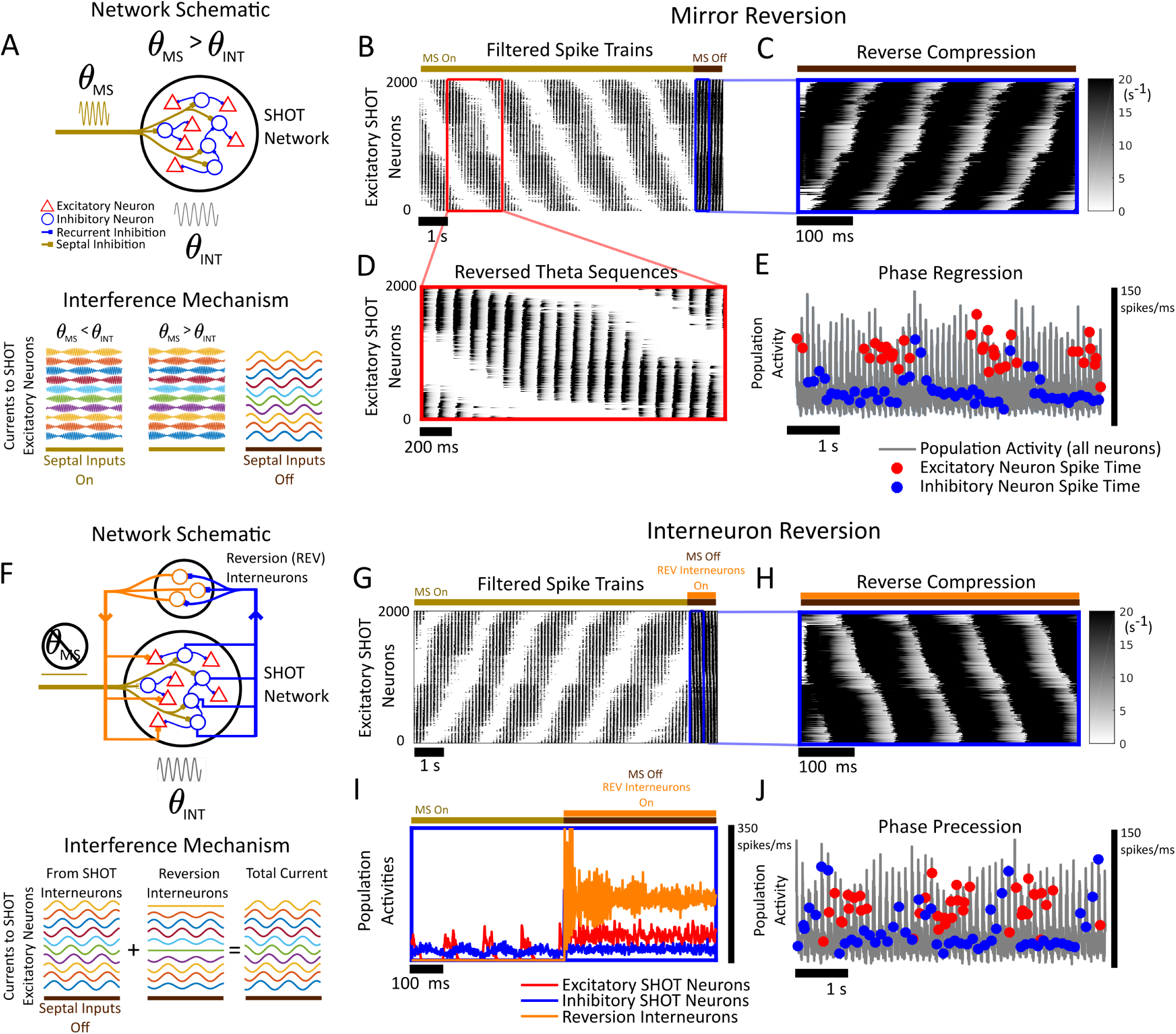
Two Mechanisms of Reverse Compression. **(A)** (Top) A network schematic of reverse compression when the septal oscillation frequency is faster than the recurrent oscillation (*θ_MS_* > *θ_INT_*). Only the septal inputs (gold) of the SHOT network are altered. (Bottom) In Mirror Reversion, interference theory predicts the reversion of time-fields when septal inputs are present (gold) relative to the order of burst firing when the septal inputs are removed (brown). **(B)** The filtered spikes (***r***(*t*), *τ_D_* = 20 ms, *τ_R_* = 2 ms) fired by the excitatory SHOT neurons. Darker values indicate the presence of spikes. The overline denotes the presence (gold, first 10 seconds) or absence (brown, last 1 second) of septal inhibition. A compressed segment is outlined in blue. The excitatory neurons (black) have time fields that occur in a decreasing sequence relative to the neuron index (higher indices fire first). **(C)** The zoomed 500 ms (highlighted in blue) segment in (B). The bursts occur in reverse order as in the time fields from (B). **(D)** A zoomed in 2 second segment of activity from (B) while septal inhibition is applied. Note that the sequence of spiking within a firing field is also reversed in order, thereby generating reversed theta sequences. **(E)** Both excitatory (red dots, spike times) and inhibitory neurons (blue dots, spike times) regress in phase relative to the network theta-oscillation (grey). The population activity is used as the theta reference. This is computed as the histogram of spike-times with 1 ms time bins. **(F)** (Top) A network schematic of Interneuron Reversion, a reverse compression mechanism using a dedicated population of interneurons to trigger reverse compression. In this mechanism, the reversion interneuron population (orange) receives non-specific and sparse connections from the inhibitory population (blue) in the SHOT network to lock the population into the recurrent theta oscillation, *θ_INT_* (grey). The reversion interneurons subsequently inhibit the excitatory neurons. If the reversion interneurons are activated when septal inhibition ceases, reverse compression occurs. (Bottom) The interference mechanism that allows for this compression is shown below the network schematic and explained in further detail in the Materials and Methods Section: Interference Based SPW-R Compression of Spike Sequences. The reversion interneurons add an additional source of inhibition to the excitatory SHOT neurons. The amplitude of this inhibition depends on the phase preference of the SHOT excitatory neurons with respect to *θ_INT_*. **(G)** The firing fields are now in the same order as in Figure 1D. The firing fields increase in order relative to the neuronal index. As in Figure 2(B), the septal inputs are on for the first 10 seconds, and deactivated for the final 1 second. Additionally, the reversion neurons are brought online at the same time. The overlines denote periods when the septal inputs are present (gold), absent (brown), and when the reversion neurons are turned on (orange). **(H)** A zoomed in, 500 ms segment of reverse compression triggered by the reversion interneurons when the septal inputs are absent. The excitatory neurons fire bursts in reverse order relative to the time-fields. **(I)** The population activity for interneurons in the SHOT network (blue), excitatory neurons in the SHOT network (red) and the reversion interneurons (orange) for 500 ms before and 500 ms after septal inhibition is removed. The reversion interneurons are brought online simultaneously to when septal inhibition ceases. The reversion interneurons display an initial synchronized transient that does not significantly affect the reversion of firing order shown in Figure 2H. **(J)** Both excitatory (red dots, spike times) and inhibitory neurons (blue dots, spike times) precess in phase relative to the network theta-oscillation (grey) when the reversion interneurons are off. The population activity is used as the theta-reference and computed as in (E).

To the best of our knowledge, phase regression has never been reported in the experimental literature while reverse theta sequences have only been reported in [Cei et al., 2014]. A second mechanism emerges if we consider external septal inputs that are slightly larger than *θ* = *θ_INT_* /2, which we term Harmonic Reversion. Like Mirror Reversion, the time-fields generated during Harmonic Reversion are in reversed order, only with phase precession instead of phase regression when *θ_MS_* < *θ_INT_* (See supplementary Figure S3). However, both Harmonic and Mirror reversion mechanisms have one critical flaw. If a memory is encoded onto the firing field backbone under Mirror or Harmonic Reversion, then the memory can only be replayed in compressed form as a reverse replay. As reverse replay is seldom seen during sleep states [Buzsáki, 2015] and primarily observed during waking states, both Harmonic and Mirror Reversion mechanisms are unlikely mechanisms of reverse replay.

Surprisingly, another mechanism for reverse compression exists if we allow for another population of interneurons. This population of interneurons both receives and projects inhibition to neurons in the SHOT network (Figure 2F-J). Thus, we refer to this mechanism as Interneuron Reversion. In Interneuron Reversion, the reversion interneurons are locked into the network theta oscillation by receiving inhibition from the SHOT interneurons (Figure 2F). The reversion interneurons then inhibit the excitatory SHOT neurons with an inhibition amplitude that depends on the phase preference of their target (see Materials and Methods, Supplementary Figure S5, Figure 2F). This is sufficient to generate reverse compression in the excitatory SHOT neurons when septal inhibition is removed simultaneous to activation of the reversion interneurons (Figure 2H-I, see Materials and Methods: Interference Based Compression of Spike Sequences, Reversion Interneuron Population). Furthermore, the phase-preference dependent inhibition need not be exact. Interneuron reversion is robust to distortions in the required relationship between the amount of inhibition an excitatory SHOT neuron receives, and its phase preference (See Supplementary Figure S5). Further, normal phase precession is observed in the presence of septal inputs when the reversion interneurons are not stimulated (Figure 2J). Thus, inhibitory interneurons can not only control and gate the timing of spikes during compression, they can reverse spike timing during compressed replay.

### 2.5 Online and Immediate Learning Using a Theta Backbone

With a robust, compressible, and reversible backbone, we investigated how these internally generated theta sequences can be used for the rapid formation and replay of episodic memories. One possible solution is to use the incoming stimulus as a supervisor and learn a set of synaptic weights in the form of decoders *ϕ*, with synaptic plasticity (Figure 3A). If the spike train is replayed, the decoder acts as the storage medium or memory engram and the memory can be read out as the product of the decoder and the repeating internally generated spike train. Thus, we derived a learning rule based on the theory of function approximation (see Supplementary Materials Section S1: The Fourier Rule). This learning rule, which we refer to as the Global Fourier rule, can be thought of as a kind of dynamic regression. The supervisor is approximated by dynamically regressing onto the internally generated theta sequences in the SHOT network, thus forming a memory engram. The rule is formulated as a differential equation:

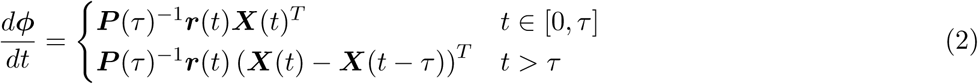

where ***r***(*t*) is the filtered spike trains from the SHOT network, ***X***(*t*) is the supervisor to be remembered, and *ϕ*(*t*) is the decoder or memory engram. The parameter *τ* serves as the period of the internally generated theta-sequence ***r***(*t*). The matrix ***P*** (*τ*) is a constant matrix that maximizes the use of the neural correlations in the SHOT network (see Supplementary Materials Section S1 for further information).

**Figure 3:**
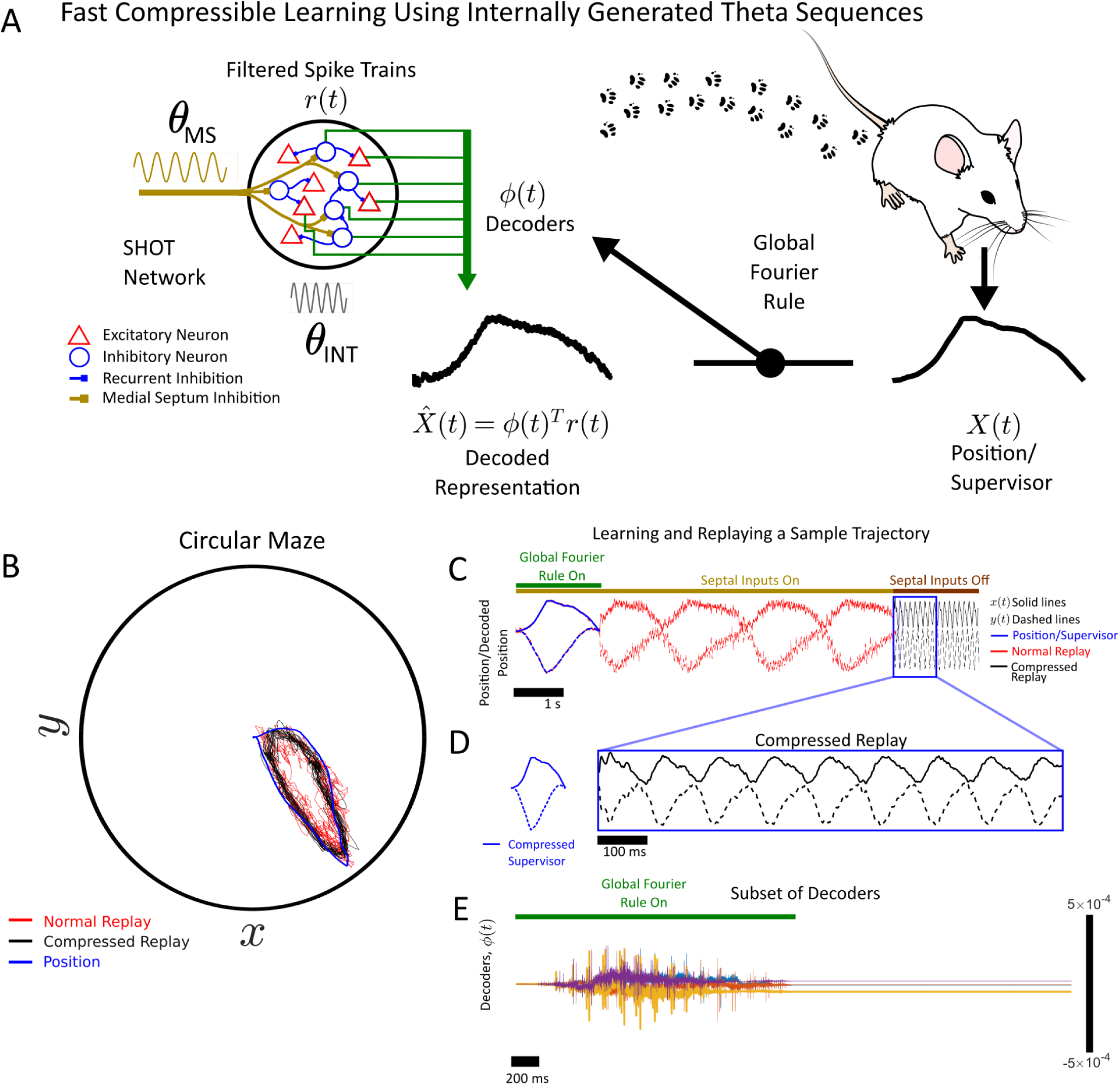
Learning Online and with a Single Stimulus Presentation. **(A)** A schematic for how decoders, *ϕ*(*t*) (green), can be learned online using the global Fourier rule applied to the SHOT network. These decoders instantly bind the supervisor or quantity to be remembered to the SHOT network activity, over the time period [*t* − *τ*, *t*] where *τ* is the period of the firing fields. As the firing fields repeat, the decoded memory replays the the decoded representation (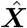 (*t*)) of the memory, (***X***(*t*) = (*x*(*t*)*, y*(*t*))). The decoder *ϕ*(*t*) in conjunction with the filtered spike trains, ***r***(*t*) for the neurons in the SHOT network, allows the network to replay its internal representation of *x*(*t*). The memory to be replayed is a simulated mouse trajectory. **(B)** A simulated mouse (blue) runs a loop around the maze. The SHOT network learns online decoders and replays the trajectory in normal time (red) or compressed time (black). **(C)** The septal inputs are present for the first 11 seconds (gold overline) and absent for the remaining 2 seconds (brown overline). The global Fourier rule is turned on for the initial 2 seconds (green overline) so that the decoders can be learned using the supervisor (blue). The memory is replayed at a normal time scale when the septal inputs are on (black, *x* component is solid, *y* is dashed) and replayed at a compressed time scale when the septal inputs are removed (black). **(D)** A zoomed in 1 second segment of the replays on a compressed time scale when the septal inputs are removed (black *x* component is solid, *y* is dashed). A compressed and time aligned version of the memory (blue) is shown for reference. **(E)** The evolution of the decoders as they are learned by the global Fourier rule (first four seconds of (C)).

The decoders are learned online and only use the stimulus once, in the initial presentation. To test this online decoder algorithm, we considered a synthetic mouse trajectory (Figure 3B) and learned a set of decoders for the SHOT network online using the mouse trajectory as a supervisor. The stimulus was successfully learned with septal inhibition present (Figure 3C). The decoders are learned immediately (Figure 3E) and with only a single presentation of the stimulus at the start of the simulation. Removal of septal inhibition triggers compression of the replay (Figure 3C-3E).

While online and immediate decoding for a network of neurons is one way of forming a memory, there are problems with this approach. First, in order to learn a memory engram online, the network requires some information about the correlations between spikes in the network via the matrix ***P*** (*τ*). This information is non-local as it depends on every neuron in the network, thereby rendering this approach biologically implausible. Second, because the spiking was strongly correlated and phase locked to the theta oscillation, we required many neurons with heterogeneity in their driving currents for this procedure to work (see Materials and Methods). Finally, it is unclear if neurons have access to the supervisor ***X***(*t*) or what the supervisor even is for that matter [Brette, 2017].

Given the problems associated with the Global Fourier learning rule, we sought alternatives that could still satisfy online and single-trial learning. First, we note that by replacing ***X***(*t*) with ***r***_*post*_(*t*), the network no longer learns a supervisor but a downstream spike train associated with a set of postsynaptic neurons. We refer to these neurons as Read Out (RO) neurons. In this case, the decoder is no longer a decoder but a synaptic weight between presynaptic excitatory SHOT neurons and postsynaptic RO neurons. Finally, as spikes are inherently discrete events that are local in time, one does not need to use the neuronal correlations optimally. This implies that ***P*** can be replaced by a constant scalar *λ* yielding:

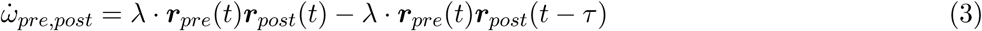

This is a Hebbian learning rule (*λ*·***r**_pre_*(*t*) ***r**_post_*(*t*)), with a forgetting term (*λ*·***r**_pre_*(*t*) ***r**_post_*(*t*−*τ*)), where *λ* dictates the rate of learning. We refer to this as the local Fourier rule.

We tested the local Fourier rule (Equation 3) with a set of RO neurons (Figure 4A-C). We can think of these neurons as being stimulated by external activity, such as if a mouse was moving along a linear track while receiving external position cues. Thus, we triggered externally evoked sequential spiking in these neurons while simultaneously presenting septal inhibition (Figure 4B,C) to the SHOT network, and running the local Fourier rule (3). The rule learns a set of weights in an online fashion between the SHOT excitatory neurons and the RO neurons. Critically, these are too weak to trigger spiking in the presence of septal inputs and do not interfere with externally induced spike trains. However, if septal inputs are removed, then compressed bursts are triggered in the SHOT network which triggers spiking in the RO neurons (Figure 4D). The spiking occurs in compressed sequential order to the spiking observed on the behavioral scale.

**Figure 4:**
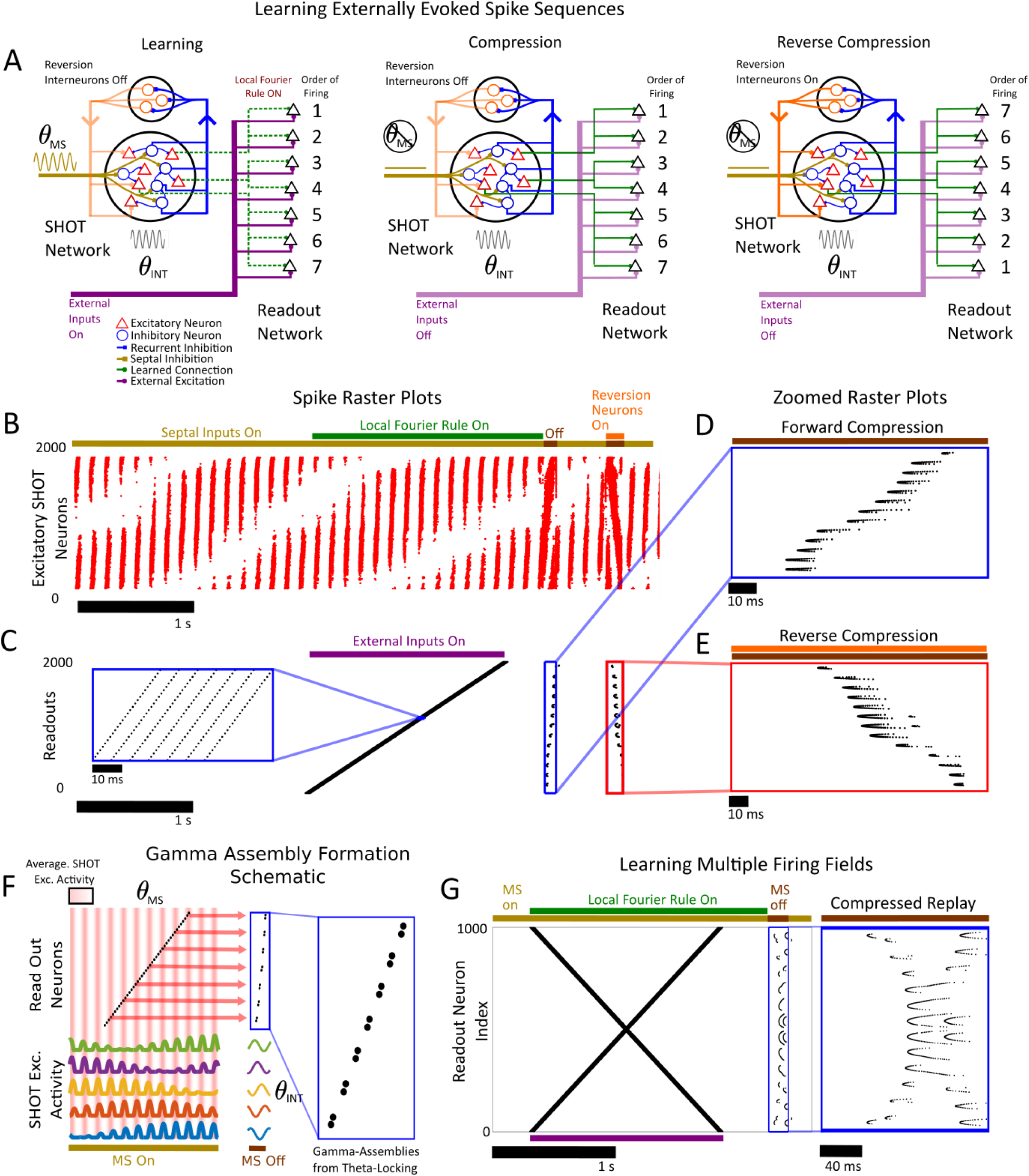
Rapid Spike Train Learning with a Septal-Hippocampal Oscillatory Theta Network. **(A)** Schematic of the network consisting of the the Septal-Hippocampal Oscillatory Theta (SHOT) network (red for excitatory, blue for inhibitory), the reversion interneurons (orange), and the readout (RO) network (black). (Left) In the learning phase, the SHOT network is inhibited by septal inputs (gold) thereby generating time-fields. A set of feedforward weights are learned between the SHOT network and the 2000 neurons in the RO network using the local Fourier rule (connections in green). The RO neurons are activated in sequential order by external inputs (in purple). (Middle) In the compression phase, both the local Fourier rule and the septal inputs are turned off, triggering compressed bursts in both the SHOT and RO neurons. (Right) Reverse compression can be elicited by activating the reversion interneurons simultaneously to removal of the septal input. **(B)** The spike-raster plot for the excitatory neurons in the SHOT network (red dots). The overlines denote the presence (gold) or absence (brown) of septal inhibition, activation of the local Fourier rule (green), and activation of the reversion neurons (orange). The septal inputs are on for the first 4 seconds during encoding and are turned off twice (*t* ∈ [4.015, 4.11], *t* ∈ [4.54, 4.675]). In the second interval, the reversion interneurons are also brought online to trigger a reverse replay. **(C)** Spike-raster plot for the excitatory neurons in RO network (black dots). The purple overline denotes the presence of external inputs triggering sequential burst firing (inset in the left is a zoom). **(D)** A zoomed in segment of forward compressed replay of the readout neurons when the septal inputs are turned off. Note that the compressed replay takes the form of a sequence of gamma assemblies. **(E)** A zoomed in segment of reverse compressed replay of the readout neurons when the septal inputs are turned off and the reversion neurons are brought online. **(F)** A schematic for the emergence of gamma assemblies from a combination of theta-phase segregation of spiking and the local Fourier rule. As the activity in the excitatory SHOT neurons is segregated to specific phases in the theta oscillation, only spikes that are coincident with that phase in the RO neurons get bound by the Fourier rule. This is due to the Hebbian nature of the Fourier rule. This creates gamma-assemblies during compression as only spikes in the RO that coincide in phase with the excitatory SHOT neurons are remembered. **(G)** (Left) Similar as in (B), only with a different downstream spike sequence (*X* shape) where neurons have multiple firing fields. Here, the septal inputs are turned off only once to trigger compressed forward replay. The reversion interneurons are not brought online. The septal inputs are turned off during *t* ∈ [3.95, 4.11] to trigger a compressed replay. (Right) A zoomed in segment of the compressed spike-replay when the septal inputs are off. Once again that the shape of the supervisor has been both compressed and discretized into a series of gamma-assemblies.

Surprisingly, this compression does not capture all spikes in the readout neurons. In fact, the local Fourier rule combined with phase precessing theta assemblies creates gamma-assembly segregation in the RO neurons during compressed replay (Figure 4F). Only RO neurons that fire in phase with the theta-locked, excitatory SHOT neurons, are remembered due to the Hebbian nature of the learning rule. This implies that theta oscillations and Hebbian learning rules can generate of gamma assemblies. The frequency of gamma-assembly firing, Γ is given by 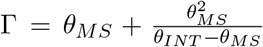 (see Materials and Methods: Gamma-Assembly Frequency as a Function of *θ_INT_* and *θ_MS_*) For the parameters we have chosen, this yields a Γ = 136 Hz. As the recurrent inhibition frequency is fixed, our result implies that septal inhibition controls both theta frequency modulation and behavioral time-scale firing fields in its presence, and gamma assemblies during compression in its absence. Further, unlike other mechanisms of gamma-frequency firing ([Whittington et al., 2000a, Börgers and Kopell, 2003, Whittington et al., 2000b, Bartos et al., 2007]) the gamma here is learned through Hebbian plasticity operating on phase precessing theta assemblies rather than fixed circuitry in down-stream circuits. Stimulating the reversion interneurons (Figure 4E) triggers a reversed replay of these gamma assemblies via Interneuron Reversion. Finally, more complicated spiking patterns can be learned (Figure 4G) with the resulting compressed replay also displaying gamma-assembly discretization. Our results imply that septal inhibition in conjunction with Hebbian plasticity can regulate spike timing on the short (gamma), intermediate (theta), and long (behavioral) time scales simultaneously.

### 2.6 Triggering Sharp-Wave Ripples With Compression

Finally, for a direct comparison to hippocampal function, we mapped these distinct populations of neurons to hippocampal anatomy. Here, we regard the SHOT network as a subset of the cells in CA3, while the RO network was a subset of cells in CA1 (although see Discussion for other possibilities). The excitatory neurons in CA3 will form synapses with excitatory neurons in CA1 via the Schaffer-Collateral Pathway and the local Fourier rule. Further, unlike previously (Figure 4), a population of interneurons was added to the RO network as our model of CA1 (Figure 5). The CA1 excitatory neurons can only excite CA1 inhibitory neurons while the CA1 inhibitory neurons can only inhibit CA1 excitatory neurons (Figure 5A). This is similar to network topologies in CA1 considered by previous researchers [Whittington et al., 2000a, Börgers and Kopell, 2003, Whittington et al., 2000b] as a mechanism for sharpwave ripple generation (see [Buzsáki, 2015] for a recent review). Other topologies, such as direct excitation from CA3 excitatory neurons onto CA1 interneurons are also possible [Donoso et al., 2018, Schlingloff et al., 2014] and can be readily incorporated in future work.

**Figure 5:**
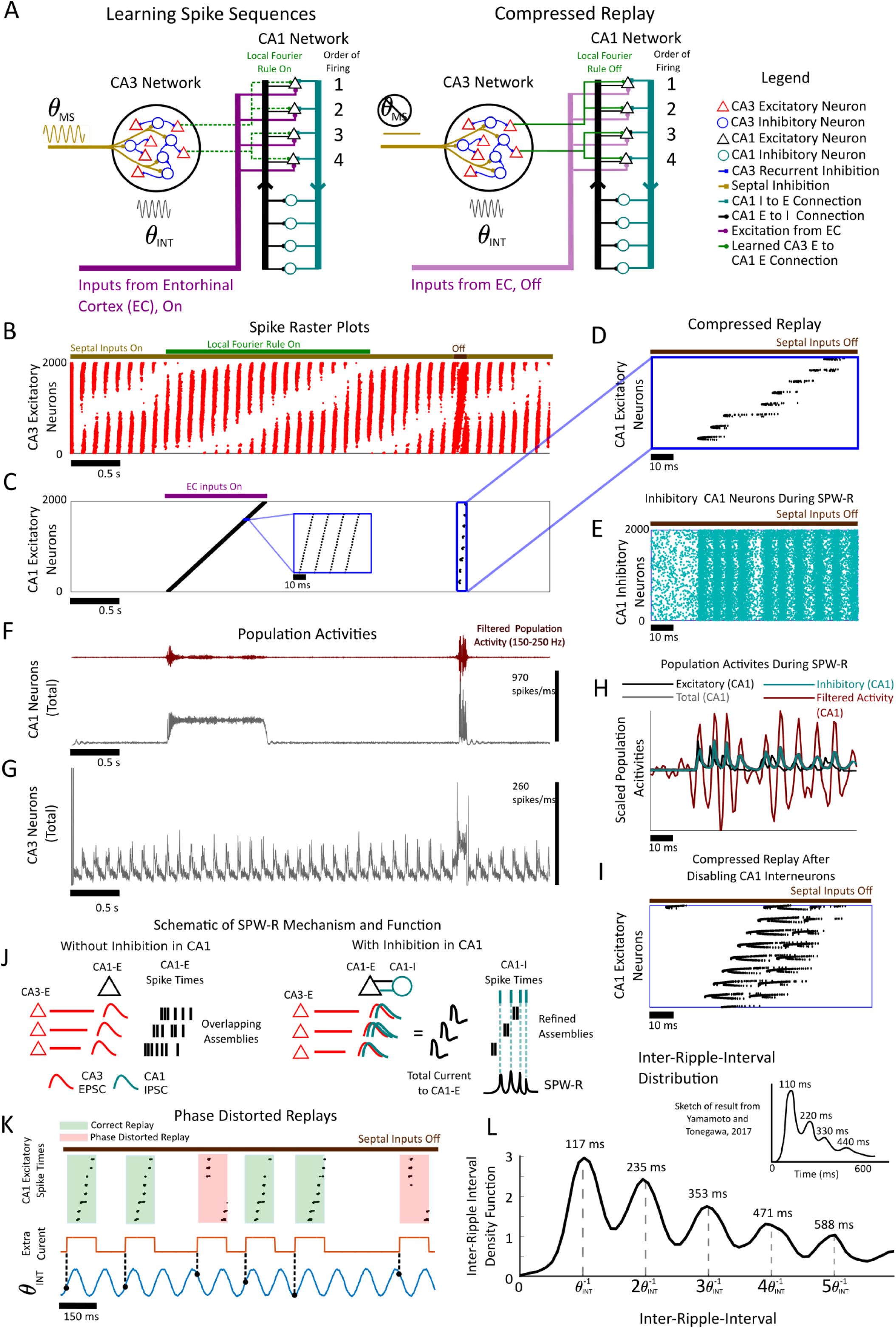
Rapid Spike Train Learning in Hippocampus. **(A)** Network schematic where the Septal-Hippocampal Oscillatory Theta (SHOT) network serves as a model for CA3, a downstream readout (RO) network serves as a model for CA1, and the external inputs are assumed to arrive from the Entorhinal Cortex (EC, purple). The CA1 network only contains *E* → *I* (black) and *I* → *E* (teal) connectivity without any *E* → *E* or *I* → *I* connections. (Left) In the learning phase, inputs from the EC trigger sequential firing in the CA1 excitatory neurons (black). Both the local Fourier rule and septal inhibition (gold) are on. The local Fourier rule binds the CA1 spiking activity to the firing fields in the CA3 excitatory neurons (red). (Right) After the connections (green) have been learned, compressed replay can be triggered by turning off septal inhibition. **(B)** The spike raster plot for excitatory neurons in the CA3 network (red dots). The overlines denote the presence (gold) or absence (brown) of septal inputs, and the activation of the local Fourier rule (green). **(C)** The spike raster plot for excitatory neurons in CA1 (black dots). The purple overline denotes the presence of inputs from EC. The CA1 excitatory neurons are stimulated to fire bursts (see inset) by EC inputs during *t* ∈ [1, 2]. The local Fourier learning rule is operating in the intervening time period and the septal inputs are on for this duration. The septal inputs are turned off during [4.03, 4.13] to trigger a compressed replay. **(D)** A zoomed segment of the raster plot in (B) showing compressed gamma assembly replay triggered when septal inhibition is off. **(E)** The spike raster plot for the CA1 interneurons (teal dots) showing high frequency firing in response to compressed replay while septal inhibition was removed. This plot is time aligned with (D). **(F)** The population activity for the CA1 population (grey). The population activity is computed as a histogram across the network with 1 ms spike times. This plot is time aligned with (B) and (C). The filtered population activity (burgundy) is band-pass filtered in the ripple band range, [150, 250] Hz. **(G)** The population activity in the CA3 region (grey). Note that for both (F) and (G), all neurons (both excitatory and inhibitory) are used to compute the population activity. **(H)** A zoomed in segment of the total CA1 population activity (grey), the inhibitory CA1 population activity (teal), the excitatory CA1 population activity (black), and the filtered ripple band (burgundy). Note that the population activities have been normalized to have the same maximum activity, and are not to scale. The majority of the population activity is due to the inhibitory neurons. The high frequency oscillations are triggered by the excitatory population which also displays gamma modulated activity. The CA1 excitatory gamma assemblies trigger the inhibitory network to fire at a gamma-frequency. **(I)** A compressed replay triggered by removing septal inputs while disabling the interneurons in CA1. The CA1 interneurons were clamped below threshold to prevent firing. This causes the learned gamma-assemblies to overlap extensively. **(J)** A schematic for sharp-wave-ripple formation and gamma-assembly refinement. The CA3 excitatory neurons trigger ordered gamma assembly spiking in CA1 through their Excitatory Post Synaptic Potentials (EPSPs). Without inhibition, the assemblies overlap extensively. When inhibitory neurons are included in CA1, the excitatory assemblies trigger the CA1 inhibitory neurons to fire. The CA3 EPSPs are refined in width by CA1 inhibitory post synaptic potentials. This refines the burst widths generated by CA1 excitatory neurons while creating SPW-R’s. **(K)** The septal currents are kept off and sharp-wave-ripples are initiated by stochastically turning on an extra current (in orange) with a constant probability per unit time (see Materials and Methods). This again triggers burst firing in the CA3 excitatory neurons. The learned connections from CA3 to CA1 excitatory neurons are identical as in (B). The extra current has a relative refractory period to model the refractory nature of sharp-wave-ripples (see Materials and Methods). Some replays are correct (green), while other replays are fragmented (red). One of the FORCE decoded *θ_INT_* oscillators (teal) demonstrates that correct replays are locked to the ascending component of the oscillation while fragmented replays are locked to the descending phase. Thus, we term this type of error phase distortion. **(L)** Phase distortion may be minimized by locking the probability of ripple initiation to the underlying *θ_INT_* oscillation in the CA3 excitatory neurons. Here, the probability of ripple initiation was now non-constant and dependent on one of the decoded oscillators of CA1 inhibitory neurons. The network was run for 3000 seconds under this condition to generate an inter-ripple-interval distribution. The inter-ripple-interval distribution is a good qualitative match for the multi-modal distribution from [Yamamoto and Tonegawa, 2017] (top inset) which has peaks at the harmonics of a theta oscillation period of 110 ms.

The CA1 excitatory neurons were (as in Figure 4A) stimulated to spike in a specific order from external inputs (Figure 5C). We interpret these inputs as arising from the Entorhinal Cortex (EC) and represent spike sequences induced by sensory stimuli that need to be remembered. Once again, the local Fourier rule was on during the presentation of septal inhibition to construct a memory engram. The septal inputs were removed and successfully triggered a compressed replay of the EC evoked spiking (Figure 5B-D). The population activity (Figure 5F) in CA1 displayed ripple-like high frequency oscillations during compression. The population activity in CA3 however demonstrated a sharp increase in activity when septal-inputs were removed (Figure 5G), reminiscent of SPW’s during compression. This is a combined effect of removing septal inhibition from CA3 inhibitory neurons, and applying a super-threshold current to CA3 excitatory neurons. Once again, compression resulted in gamma-assembly formation in the CA1 excitatory neurons (Figure 5C,D) while the CA1 inhibitory neurons fired synchronous volleys at a gamma-frequency (Figure 5E). The number of gamma assemblies in the excitatory neurons was less than the number of bursts triggered in the inhibitory population (8 vs. 10). In general, depending on how strongly CA1 excitatory neurons excite the CA1 inhibitory neurons, higher ripple frequency oscillations can emerge.

A closer look at the population activity of the excitatory and inhibitory population reveals that the majority of the population activity is due to the inhibitory neuron population in CA1 as the inhibitory neurons were locked to the population activity. If we regard the population activity as a proxy (albeit a poor one) for the local-field potential, than our results mirror PV basket cells locking to the LFP ripples [Ylinen et al., 1995, Klausberger and Somogyi, 2008, Klausberger et al., 2003, Klausberger et al., 2004, Klausberger et al., 2005]. However, we caution that this result should be investigated further by using generative LFP models in the future (see [Mazzoni et al., 2015, Chatzikalymniou and Skinner, 2018] for example). The CA1 excitatory neurons also displays some locking to the ripples (Figure 5H). In fact, the gamma-assemblies precede the ripples and are indeed causal. As the CA1 excitatory cells synapse with random and all-to-all coupling to the inhibitory population, they trigger bursts in the inhibitory population at a gamma-frequency. This is similar to the Pyramidal Interneuron Network Gamma (PING) mechanism operating with external sequential driving of the excitatory CA1 neurons [Whittington et al., 2000a, Börgers and Kopell, 2003] or the feedback inhibition/PYR-INT model [Buzsáki, 2015, Whittington et al., 2000b]. We also note that many models of ripple generation exist ([Whittington et al., 2000b, Brunel and Wang, 2003, Taxidis et al., 2012, Taxidis et al., 2013, Donoso et al., 2018], to name a few) and we leave an analysis of how compressed replay triggered by CA3 interacts with these different ripple generation mechanisms for future work.

As the SHOT and Reversion interneurons coordinated spike timing, we investigated if the CA1 inhibitory neurons performed a similar function. To test this hypothesis, we disabled the interneurons post-learning while triggering a compressed replay (Figure 5I) by removing septal inputs. The gamma-assemblies triggered by the neurons were no longer segregated to discrete moments in time and overlapped extensively. In fact, the first gamma-assembly persisted and overlapped with the last gamma-assembly during the replay. This implies that the inhibitory neurons in the CA1 layer might be refining the duration of the learned gamma assemblies and segregating them to their own discrete units of time (Figure 5J). This may be an advantageous strategy to transmit information reliably out of the hippocampus. For example, CA1 interneurons prevent spikes from non-sequential assemblies from additively summing and triggering temporally imprecise spikes in downstream cortical networks. Alternatively, this discretization may be advantageous for spike-timing-dependent plasticity in downstream networks. As with the SHOT and Reversion interneurons, our results demonstrate that interneurons in CA1 also control the timing of spikes by refining the width of learned gamma assemblies in CA1 via a PING-like mechanism [Whittington et al., 2000a, Börgers and Kopell, 2003].

### 2.7 Gamma Assemblies Nested on a Theta Oscillation

Gamma assembly discretization occurs during sharp-wave-ripples in our model. However, gamma oscillations are often nested with theta oscillations and implicated in the formation of memories [Belluscio et al., 2012, Lisman, 2005, Heusser et al., 2016, Lisman and Buzsáki, 2008, Tukker et al., 2007]. Further, gamma oscillations are found in both theta and non-theta states (during SPW-R’s) [Buzsáki, 2015, Tukker et al., 2007, Csicsvari et al., 2003]. Thus, we investigated if these gamma-assemblies would persist in the presence of septal inhibition (Supplementary Figure S6). If the CA3 excitatory neurons are provided with slightly more excitation in the presence of septal inputs, the CA1 excitatory neurons fire as gamma assemblies nested on a theta oscillation. This occurs after learning with the local Fourier rule (as in Figure 5). Thus, theta phase segregation and simple Hebbian learning rules can generate theta-nested gamma assemblies that function similarly to a working memory buffer first suggested in [Lisman and Idiart, 1995, Jensen and Lisman, 1996]. Here, however, the maximum number of discrete items, *N_items_*, that can be stored in the buffer is dictated by (*θ_MS_*)^−1^ and (*θ_INT_* − *θ_MS_*)^−1^:

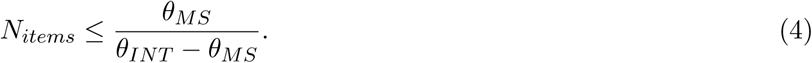

The period of the septal oscillation (*θ_MS_*)^−1^ acts as the temporal resolution while the time cell periodicity (*θ_INT_* − *θ_MS_*)^−1^ determines length of the buffer (Materials and Methods: Gamma-Assembly Frequency as a Function of *θ_INT_* and *θ_MS_*). For the parameters we have chosen, we arrive at a maximum of 16 items in the buffer.

### 2.8 Ripple-Clusters Minimize Fragmentation of Memories

Finally, we investigated how the supposed stochastic nature of SPW-R initiation would alter replays [Buzsáki, 2015, Schlingloff et al., 2014]. Previously, we disabled septal inputs and activated the excitatory CA3 neurons with external excitation at specific times to trigger compressed replay in CA1. To account for the potentially random nature of SPW-R initiation, we disabled the septal inputs post-learning and stochastically turned on extra excitation to initiate sharp-wave-ripples (Figure 5K) in lieu of precisely timed excitation. This is a phenomenological model of the stochastic initiation of ripples and caused stochastic replays of the learned memory (seen in Figure 4B). We found that depending on when the ripple was initiated, replays could display an error which we term phase distortion. In phase distortion, the replay is fragmented into pieces due to the underlying *θ_INT_* oscillation in the CA3 inhibitory neurons. Each fragmented segment replays a small component of the memory, however the fragments are disordered. Phase distortion occurs when the SPW-R’s are initiated over a phase range in the *θ_INT_* oscillation (Figure 5K). Thus, to reduce phase distortion errors, the inhibitory CA3 neurons can bias the probability of ripple initiation to the correct phase range of the CA3 oscillation *θ_INT_*. We considered the probability *p*(*t*) of initiating a SPW-R as

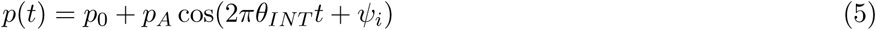

where *ψ_i_* is the phase that minimizes the phase distortion error. The parameter *p*_0_ serves as the background ripple activation rate and *p_A_* determines how strong the ripple generation probability oscillates. The oscillatory component is decoded from the CA3 inhibitory neurons (see Materials and Methods). A relative refractory period was also incorporated such that the probability of initiating immediately successive ripples is reduced (see Materials and Methods). This phase biasing inhibition has an observable effect on the inter-ripple-interval. Indeed, the inter-ripple-interval displayed a strong *θ_INT_* modulation due to this inhibitory bias (Figure 5L). This displays a strong similarity to the inter-ripple-intervals recently reported in [Yamamoto and Tonegawa, 2017]. The inter-ripple-intervals from [Yamamoto and Tonegawa, 2017] are multi-modal with peaks at 110 ms, 220 ms, 330 ms, and 440 ms for mice navigating large mazes. The interval results from [Yamamoto and Tonegawa, 2017] imply some theta modulation in ripple-genesis with a potential *θ_INT_* ≈ 9.1 Hz. Thus, our results demonstrate how inhibition can not only control spike timing within SPW-R’s, but influence SPW-R initiation and inter-ripple-interval distributions to minimize the fragmentation of memories.

## 3 Discussion

Our memory systems have to handle multiple simultaneous constraints in order to record, store, and catalogue important events. The most fundamental of these constraints is that we can only guarantee a single stimulus presentation, as it is happening. Thus, we investigated whether it was possible to bind external stimuli or the spike trains generated by the environment onto background spiking activity. The background spiking activity took the form of internally generated theta sequences or time cells. These sequences were generated through an interference mechanism involving an inhibition mediated, intrinsic theta oscillation. The second theta-frequency was externally applied and corresponded to inhibition from the medial septum. We found that this was sufficient to create stable firing fields through an interference pattern generated in the excitatory neurons. Removal of septal inhibition can trigger compression of the time-fields as subthreshold oscillations in the excitatory neurons in the SHOT network. The sub-threshold oscillations had phases dictated by the order of time-fields in the presence of septal inputs. Ordered and compressed bursts could be initiated in the excitatory SHOT neurons by applying either an external current, using dense recurrent excitation, or disinhibition through an increase in septal inhibition. The mathematics behind interference theory demonstrated three possible mechanisms for reverse replay. Of the three mechanisms, Interneuron Reversion was the most plausible and required another population of interneurons.

After having constructed this compressible temporal backbone, we derived a local learning rule that exploited the underlying oscillations in the network for fast and single-instance learning. Our learning rule could successfully bind spike-trains that were generated by external inputs to the internally generated sequence in the SHOT network immediately. Triggering a compression in forward or reverse modes would trigger a compressed forward or reversed replay. Surprisingly, through a combination of the local Fourier rule and theta frequency segregation during encoding, gamma-assemblies emerged during compressed replay. These gamma assemblies also persisted in the presence of septal inhibition as gamma assemblies nested on a theta oscillation. Finally, we found that coupling downstream networks modelling CA1 with the SHOT network modelling CA3 triggered high-frequency oscillations similar to sharp-wave ripples. The gamma-assemblies formed during compression initiated high frequency synchronized firing in the CA1 inhibitory population. The inhibitory population of CA1 was the dominant component of the population activity, and displayed strong phase locking to the ripple-like oscillations in the population activity. Furthermore, we found that the CA1 inhibitory population served to segregate learned gamma-assemblies and prevent assembly overlap. Finally, we demonstrated how inhibition in CA3 networks can prevent memory fragmentation due to phase distortions during compressed replay by modulating the probability of ripple initiation. We predicted that this would have an observable effect on inter-ripple-interval distributions and has been recently verified experimentally [Yamamoto and Tonegawa, 2017]. Our results demonstrated the critical nature of inhibitory subpopulations to provide a backbone for stable learning. We provide theoretical evidence for how inhibitory populations can generate, compress, reverse, and refine the timing of spike sequences throughout the hippocampus to allow for one-shot, online learning.

### A Link Between Theta-Phase Compression and Sharpwave Ripple Compression

Our results demonstrate that the mechanism for sharpwave-ripple compression and theta-phase precession may be one and the same. The timing of spikes during both events can be coordinated by a single population of interneurons. The difference in these two cases is the presence of septal inhibition. When septal inhibition is present, the theta phase compression that takes place are essentially miniaturized sharp-wave ripples at the population level. When septal inhibition is removed, the phase precessing assemblies become bursts in the presence of additional excitation or a potentially axo-axonic disinhibitory mechanism [Viney et al., 2013, Somogyi et al., 2014]. In fact, this hypothesis that these two modes of compression are related is supported by three lines of experimental evidence. First, the compression ratios for theta-phase compression and SPW-R compression are similar, with SPW-R offering only a 30% improvement in compression [Buzsáki, 2015, Diba and Buzsáki, 2007]. Second, the boundary between a proper sharp-wave-ripple and normal firing is arbitrarily defined [Buzsáki, 2015]. In our work, we can continuously transition between phase-precessing time-cells to SPW-R compression by decreasing septal input amplitude while simultaneously increasing the background current to the excitatory neurons. Third, in extended replay, ripples often occur in bursts or clusters containing multiple sharpwave ripples [Davidson et al., 2009, Yamamoto and Tonegawa, 2017]. The individual ripples tend to be separated by an interval of approximately 100-150 ms, well within the theta period range [Davidson et al., 2009, Yamamoto and Tonegawa, 2017, Buzsáki, 2015]. Strikingly, in [Yamamoto and Tonegawa, 2017], the authors find theta-harmonics in the inter-ripple-interval in the form of peaks in the distributions at 110, 220, 330, and 440 ms. Here, we demonstrated that a similar distribution would arise if inhibition modulated the probability of ripple genesis as a mechanism to reduce phase distortion.

Furthermore, the SPW-R/theta dichotomy displayed by the hippocampus is naturally explained by our model as an interference between two-theta oscillations. One theta oscillation is intrinsic and inhibition mediated. This oscillation controls the timing of spikes during sharp wave ripples. The other theta oscillation is supplied by inhibition from the medial septum and initiates learning by generating behavioral time scale spike sequences to bind external information to. This immediately implies that some theta-frequency modulation occurs in inter-ripple-interval distributions which was demonstrated experimentally in [Yamamoto and Tonegawa, 2017]. We hypothesize that this is due to the interneuron population in CA3 which oscillates at a frequency of *θ_INT_* and generates the compressed backbone of spiking. Finally, the current paradigm for sharpwave genesis is that the recurrent collateral system in CA3 generates enough excitation to stochastically trigger a sharpwave [Buzsáki, 2015]. Our model does not refute this but instead conjectures that excitatory neurons with some bias from the inhibitory neurons decide stochastically when a SPW-R occurs, while the inhibitory neurons are in constant control of the spike sequence that occurs during a ripple.

### The Intrahippocampal Theta Oscillation

Multiple lines of experimental evidence suggest that the hippocampus can act as its own intrinsic theta-oscillator, and the existence of multiple-theta oscillations operating within the hippocampus. First, the intrinsic theta rhythmic generation capabilities of the hippocampus is well established in the literature [Buzsáki, 2002, Kramis et al., 1975]. In fact, two potential mechanisms are supported by experimental evidence. The first mechanism involves the CA3 recurrent collateral system [Buzsáki, 2002, Bragin et al., 1995] which may act as an intrahippocampal theta oscillator when acetylcholine is present [Konopacki et al., 1987, Bland et al., 1988]. This mechanism does appear to depend on the interneuron population [Buzsáki, 2002, Traub et al., 1992, MAcVICAR and Tse, 1989, Fellous and Sejnowski, 2000]. Alternatively, CA1 may act as an intrahipppocampal oscillator [Goutagny et al., 2009, Amilhon et al., 2015]. In which case, the CA3 region may inherit the theta oscillation induced by backprojecting interneurons in CA1 [Buzsáki, 2002]. Recent theoretical work has also highlighted how interneurons can contribute or indeed generate this CA1 based intrahippocampal theta oscillation [Ferguson et al., 2017, Ferguson et al., 2015, Sekulić and Skinner, 2017, Bezaire et al., 2016, Wulff et al., 2009]. These results imply that our hypothesized SHOT network may consist of an extended CA1 and CA3 network of neurons with the SHOT interneurons potentially located in CA1. Thus, both theoretical and experimental studies suggest that a network of interneurons can generate intrinsic theta oscillations, and multiple theta rhythms are present in hippocampal networks.

### Septal Drive

Recent work has demonstrated the potential for optogenetic deactivation and driving of the medial septum in rodents [Boyce et al., 2016, Bender et al., 2015, Vandecasteele et al., 2014, Mamad et al., 2015]. Here, we suggest a novel experiment to assess whether interference theory is indeed a mechanism for the generation of intrinsically generated theta oscillations. First, we suggest employing the left/right alternation task as in [Pastalkova et al., 2008]. Then, stable-firing fields should emerge during wheel running [Pastalkova et al., 2008, Wang et al., 2015] (Figure 6A). During wheel running, we suggest using a closed loop feedback system to optogenetically drive the medial septum to oscillate at different frequencies, as in [Bender et al., 2015] (Figure 6A). As the driving frequency is swept inside and outside the theta range, interference theory predicts that periodicity of the firing fields to vary, depending on the relationship between the driving frequency and the intrinsic theta oscillation, For large enough driving frequencies, reversions of the firing fields and theta sequences should emerge (Figure 6A). However, we caution that this can also alter the mouse running speed ([Bender et al., 2015]) which may alter the firing field periodicity. A similar experiment has been conducted only in a non-episodic memory task [Zutshi et al., 2018]. Here, the authors find that the phase precession code is not retained at optogenetic driving of the medial septum at either 8 Hz or 10 Hz frequencies while place-fields are.

**Figure 6:**
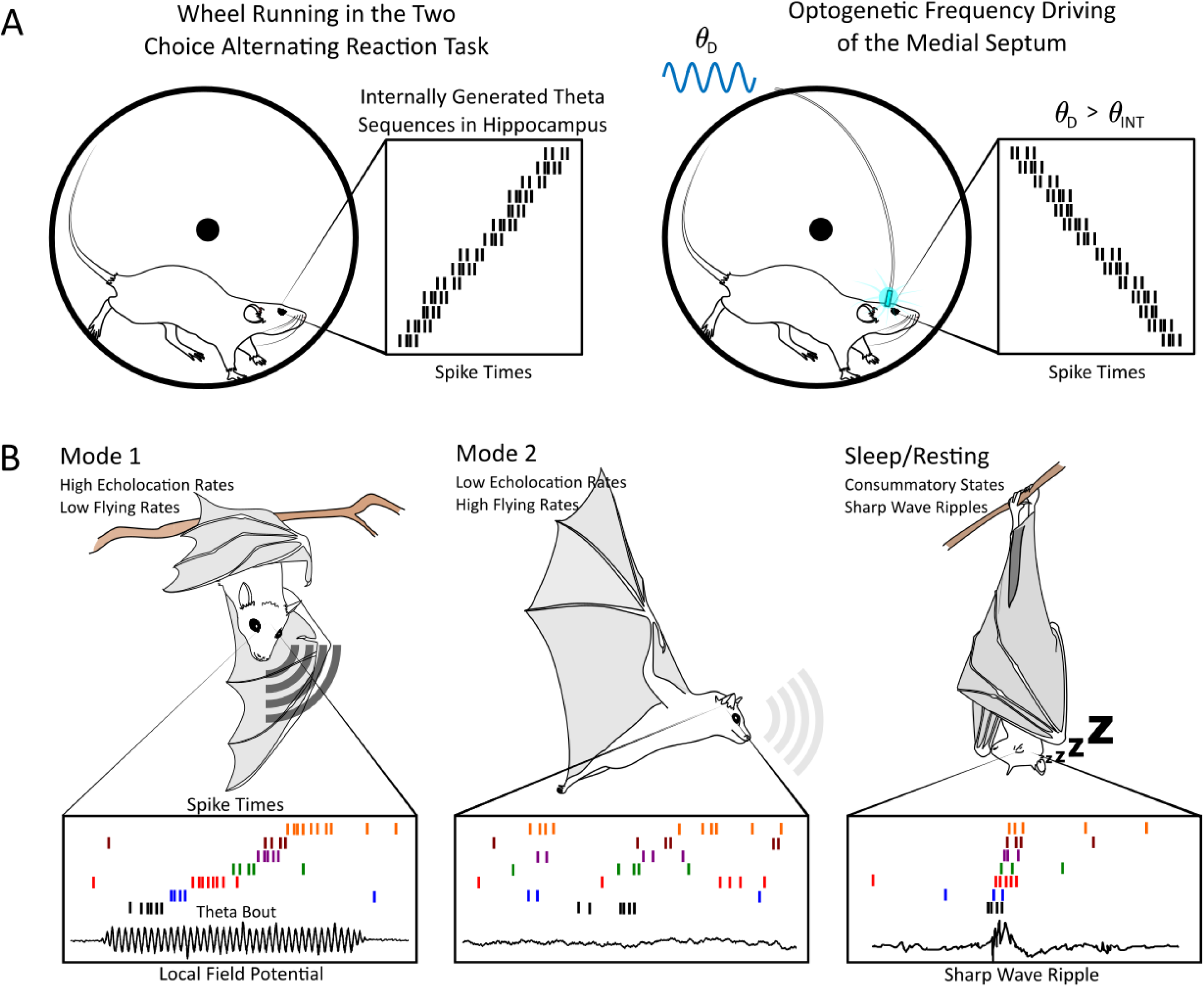
Experimental Predictions in Rodents and Bats. **(A)** Suggestion of a novel experiment to test our model of firing field formation in rodents. (Left) During the normal two choice alternating reaction task, ordered firing fields emerge during wheel running [Pastalkova et al., 2008]. (Right) We suggest running the same task, only optogenetically stimulating interneurons in the medial septum with an oscillatory input with a driving frequency of *θ_D_*. The driving frequency *θ_D_* is varied from wheel running trial to trial. When *θ_D_* > *θ_INT_*, the time-field orientation should reverse. Further, we predict that the resulting firing fields will display similar transitions as in our sweep protocol in Supplementary Figure S3. **(B)** The Egyptian Fruit Bat has two distinct behavioral modes [Ulanovsky and Moss, 2007]. (Left) In Mode 1, the bat has low flying rates and high echolocation rates. This mode also has an abundance of theta bouts in the local field potential (LFP). Theta bouts correspond to theta oscillations in the LFP that are 1-2 seconds in duration. (Middle) In Mode 2, the bat has high flying rates and low echolocation rates. Mode 2 contains less bouts of theta oscillations. (Right) We predict that compressed spike sequences that are expected to occur during sharp-wave ripples are compressed forms of the sequences observed during theta bouts, which primarily occur in Mode 1.

### Theta Oscillations in Bats

Interference Theory has been previously applied to explain how grid cells are formed [Burgess, 2008, Jeewajee et al., 2008, Koenig et al., 2011, Zilli et al., 2009, Hasselmo, 2008, Bush and Burgess, 2014, Giocomo et al., 2011]. However, recent experimental work in bats casts doubt on the role of interference models as a generative mechanism for grid cells. For example, [Yartsev et al., 2011] find grid cells in bats without the continuous theta-oscillation present in rodents. Place cells without continuous theta-oscillations are also present in bats [Yartsev and Ulanovsky, 2013]. While continuous theta-oscillations are not present in datasets involving bats, bouts of theta oscillations that are approximately 1-2 seconds occur [Yartsev et al., 2011, Yartsev and Ulanovsky, 2013, Ulanovsky and Moss, 2007].

Here, we hypothesize that these theta-bouts correspond to encoding and learning of incoming stimuli. Furthermore, we hypothesize that they are due to internally generated theta assemblies triggered by septal inhibition. In [Ulanovsky and Moss, 2007], the authors find theta-bouts that occur almost exclusively when the animal is not moving, and using echolocation to explore the environment. The authors refer to this as Mode 1. Mode 2 stands in contrast to Mode 1 and occurs when the animal is exploring the environment by moving, and using lower echolocation call rates [Ulanovsky and Moss, 2007] (Figure 6B). No clear theta peak in the local-field potential was reported in Mode 2. We conjecture that information gleaned from echolocation may be bound to internally generated theta sequences for subsequent navigation during Mode 1.

As in the bat literature, [Wang et al., 2015] also find that place cells can also exist without a background theta-oscillation by injecting muscimol into the medial septum of rats. The authors find that previously formed firing fields are still present in the absence of theta oscillations. These fields are likely driven by sensory cues in the environment, unlike internally generated theta sequences. New fields however do not form in novel mazes under muscimol injections. Further, in vivo intracellular recordings in mice navigating a virtual maze demonstrate that for both grid cells and place cells, place firing is more correlated with an underlying asymmetric ramp [Harvey et al., 2009, Domnisoru et al., 2013] rather than with an amplitude modulated theta oscillation. However, phase precessing theta oscillations with an amplitude that increases as the animal traverses through a firing field are still found in the subthreshold dynamics [Harvey et al., 2009, Domnisoru et al., 2013]. The amplitude modulations in the theta oscillations were more correlated to the timing of spikes, rather than place.

Our results lead to the conjecture that the absence of theta in grid cells in bats and the absence of the theta in place cells in rodents under muscimol inactivation are related. The theta oscillations that occur in bats and mice can be used for the purpose of learning by binding externally evoked spike trains with internally generated spike trains for later consolidation through compression. The firing fields that emerge after may be due to sharp-wave-ripple mediated construction of location dependent firing fields within the hippocampus or, attractor based dynamics that are either intrinsic, or modified by SPW-R mediated compression. Our model suggests that the theta-bouts are also the critical components for learning in bats. A similar hypothesis was put forward in [Ulanovsky and Moss, 2007]. One means of testing this hypothesis would be to compare the spiking sequences during theta-bouts (in behavioral Mode 1), sharp-wave-ripples, and sequences during behavioral Mode 2 in bats. We predict that the spike-sequences in sharp-wave ripples are compressed versions of the sequences that occur during the theta-bout intervals in Mode 1, as opposed to sequences that occur in Mode 2 (Figure 6B).

### Conclusion

A long-standing hypothesis is that interneurons control and gate the timing of spiking. Here, we demonstrate a concrete mechanism for how different populations of interneurons can 1) generate the theta-oscillation in the hippocampus through recurrent inhibition and septal inhibition 2) create phase precessing assemblies of internally generated theta sequences, 3) trigger compressed replay of these assemblies, 4) reverse a compressed replay, 5) facilitate single-instance learning of spike trains as compressed gamma-assemblies and 6) refine said gamma-assemblies by minimizing their overlap in time, 7) nest these assemblies on a background theta-oscillation and finally 8) modulate the inter-ripple-interval distribution to prevent the replay of fragmented memories. This intricate control over spiking in our model was facilitated little more than three mechanisms: Interference Theory, and Interneuron/Pyramidal reciprocal connectivity, and Hebbian plasticity. We hope that future experimental testing using optogenetic stimulation of the medial septum will test whether the intrahippocampal theta oscillator and the septal oscillator interact to generate both theta oscillations and sharp-wave ripples.

## 4 Methods

### 4.1 Leaky Integrate and Fire Network

The Septal Hippocampal Oscillatory Theta (SHOT) network consist of coupled leaky integrate-and-fire neurons:

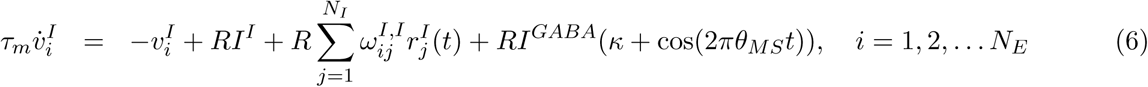

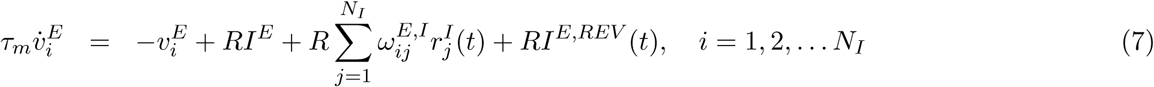

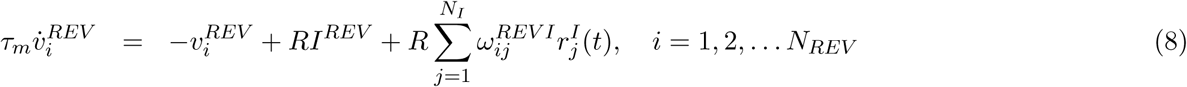

where *E*, *I* denote the excitatory and inhibitory populations of the SHOT network, respectively while REV denotes the reversion interneurons (see Materials and Methods Section 4.8). The neurons receive a constant background current *I^α^* for *α* = *E*, *I*, *REV* current set at or above the threshold (see Table 1 for parameters). The parameter *R* = 1 · 10^9^ Ω serves as the resistance. When the voltage variables reach a threshold value, *v_threshold_*, they are immediately reset to *v_reset_* followed by an absolute refractory period, *τ_ref_* during which the neuronal dynamics are quenched at the reset value. The membrane time constant, *τ_m_* determines the degree to which the synaptic currents are filtered by the voltage dynamics. The spikes themselves are filtered by the double exponential synapse:

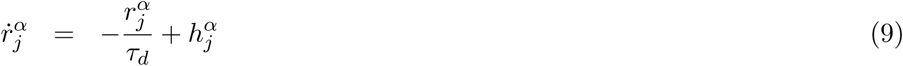

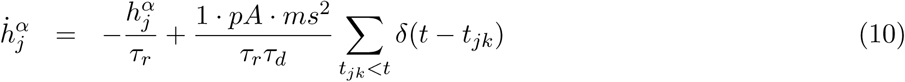

where *τ_r_* is the synaptic rise time, and *τ_d_* is the synaptic decay time *t_jk_* is the *k*th spike fired by the *j*th neuron. The inhibitory neurons in the SHOT network all receive oscillatory septal inhibition where *θ_MS_* is the input frequency, and *κ* > 1 determines the tonic level of inhibitory drive. The septal inhibition has amplitude *I^GABA^* = −10 pA for *i* = 1, 2, … *N_I_*. The excitatory neurons and REV interneurons do not receive direct septal inhibition.

**Table 1:**
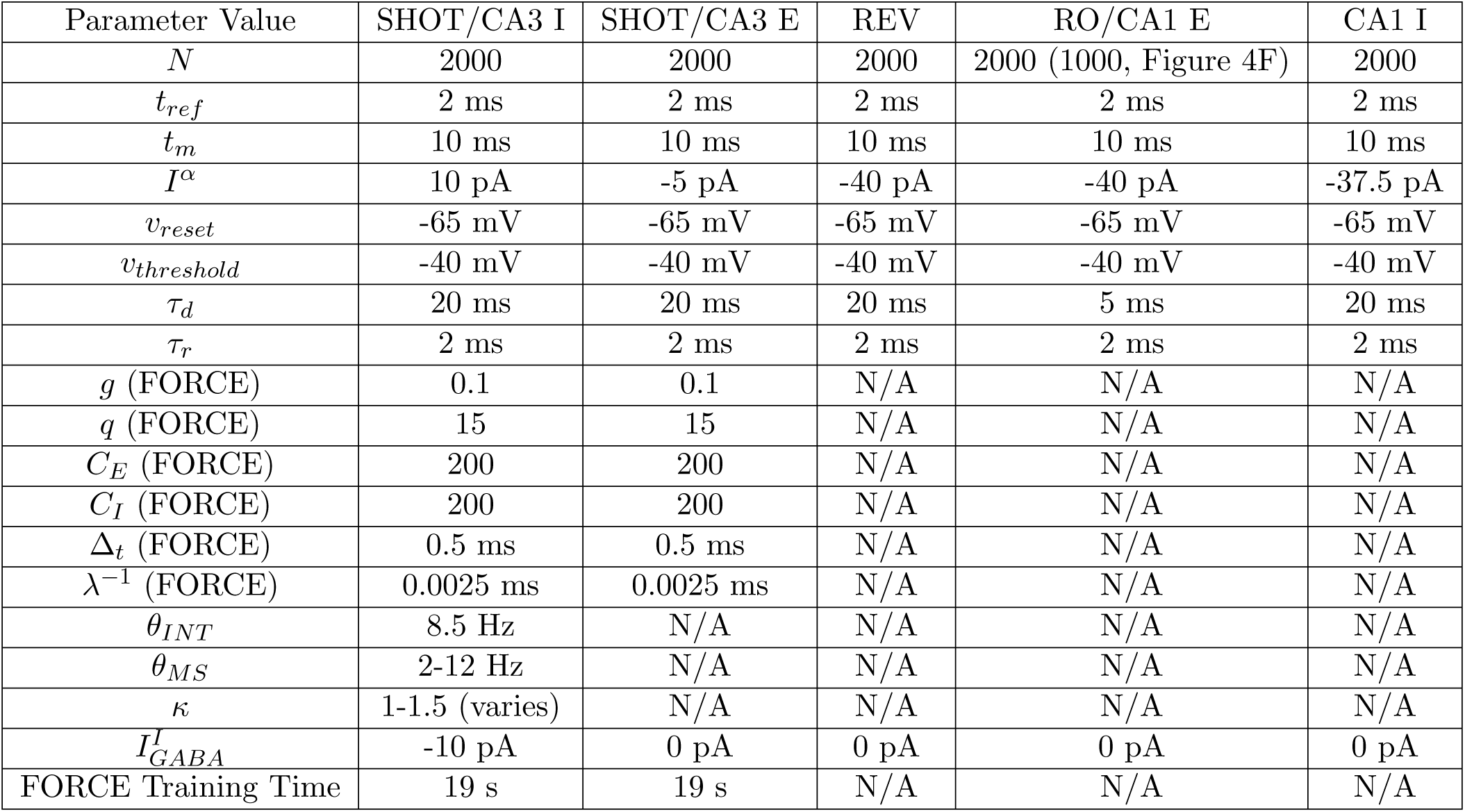
The parameters used for the SHOT/CA3 neurons, the reversion interneurons, and CA1/RO excitatory and inhibitory populations. For all networks, we use an integration time step of *dt* = 0.05 ms and Euler integration. Some of the parameters are not applicable (N/A) to all neuron models. Also note that the background current to neurons can also vary depending on the presence/absence of septal inputs. See Further Methods for Figures for more details.

| Parameter Value | SHOT/CA3 I | SHOT/CA3 E | REV | RO/CA1 E | CA1 I |
| --- | --- | --- | --- | --- | --- |
| $N$ | 2000 | 2000 | 2000 | 2000 (1000, Figure 4F) | 2000 |
| $t_{ref}$ | 2 ms | 2 ms | 2 ms | 2 ms | 2 ms |
| $t_m$ | 10 ms | 10 ms | 10 ms | 10 ms | 10 ms |
| $I^\alpha$ | 10 pA | -5 pA | -40 pA | -40 pA | -37.5 pA |
| $v_{reset}$ | -65 mV | -65 mV | -65 mV | -65 mV | -65 mV |
| $v_{threshold}$ | -40 mV | -40 mV | -40 mV | -40 mV | -40 mV |
| $\tau_d$ | 20 ms | 20 ms | 20 ms | 5 ms | 20 ms |
| $\tau_r$ | 2 ms | 2 ms | 2 ms | 2 ms | 2 ms |
| $g$ (FORCE) | 0.1 | 0.1 | N/A | N/A | N/A |
| $q$ (FORCE) | 15 | 15 | N/A | N/A | N/A |
| $C_E$ (FORCE) | 200 | 200 | N/A | N/A | N/A |
| $C_I$ (FORCE) | 200 | 200 | N/A | N/A | N/A |
| $\Delta_t$ (FORCE) | 0.5 ms | 0.5 ms | N/A | N/A | N/A |
| $\lambda^{-1}$ (FORCE) | 0.0025 ms | 0.0025 ms | N/A | N/A | N/A |
| $\theta_{INT}$ | 8.5 Hz | N/A | N/A | N/A | N/A |
| $\theta_{MS}$ | 2-12 Hz | N/A | N/A | N/A | N/A |
| $\kappa$ | 1-1.5 (varies) | N/A | N/A | N/A | N/A |
| $I_{GABA}^I$ | -10 pA | 0 pA | 0 pA | 0 pA | 0 pA |
| FORCE Training Time | 19 s | 19 s | N/A | N/A | N/A |

The weight matrices ***ω***^*I*,*I*^, ***ω***^*E*,*I*^ are described in further detail in the section FORCE Training Internal Sequences. All weight matrices that we consider are unitless with the units of current (pA) carried by the synaptically filtered spike trains *r*(*t*) (see equation (10)). The excitatory neurons also receive a current from the reversion interneuron population (in Figures 2 and 4), *I^E,REV^* (*t*) and is descried in greater detail in the section Reversion Interneuron Population. In all cases, when we refer to the filtered spike train, we mean ***r***(*t*) as given by equation (9). However, the filtering time constants used differ for different figures and different populations of neurons (see Table of Parameters). Note that the only connections in equations (6)-(8) are from the SHOT interneurons (*r^I^*(*t*)) or the reversion interneurons (*r^REV^* (*t*)).

For Figure 4, a set of readout neurons (RO) was also included in the simulation:

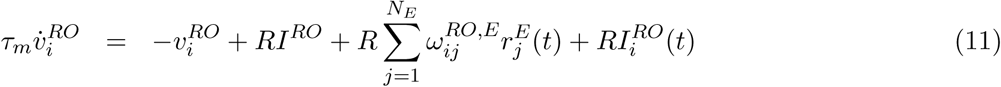

where the weights 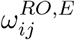 were trained using the local Fourier Rule. A series of external input currents *I^RO^*(*t*) were transiently applied to elicit bursting, and are described in the section Further Methods for Figure 4.

Finally, to model CA1 in Figure 5, we considered another population of interneurons and excitatory neurons:

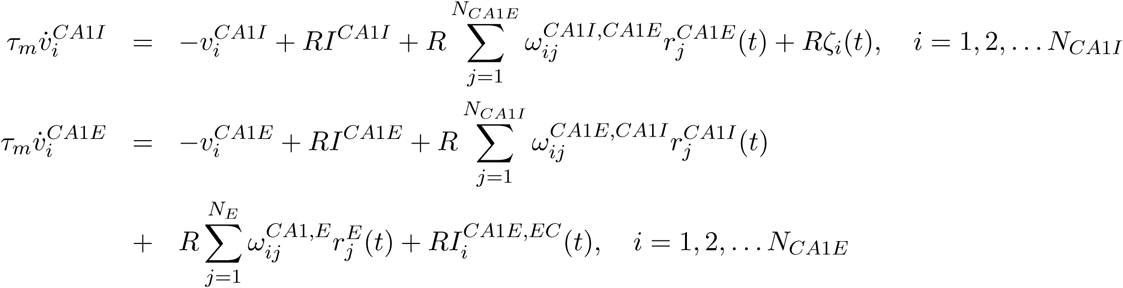

where *ζ_i_*(*t*) is an independent white noise term with mean 0 and standard deviation *σ* = 0.2 pA. The weight matrices ***ω***^*CA*1*E,CA*1*I*^, ***ω***^*CA*1*E,CA*1*I*^ are untrained and discussed further in the section Further Methods for Figure 5. The inputs from the entorhinal cortex to the CA1 excitatory neurons, 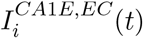 are also described in the section Further Methods for Figure 5. Finally, the weight matrix ***ω***^*CA*1*E,E*^ is trained by the local Fourier rule, which is discussed further in the Supplementary Materials Section S1. The SHOT network serves as our model of CA3 (equations (6)-(8)).

### 4.2 FORCE Training a Balanced E/I Network with Dales Law

The weight matrices ***ω***^*αβ*^ = ***S***^*αβ*^ + ***L***^*αβ*^ are decomposed into static terms (***S***^*αβ*^) that are chosen to form an initial balanced spiking network, and learned terms (***L***^*αβ*^) that are FORCE trained [Nicola and Clopath, 2017]. The static term is given by:

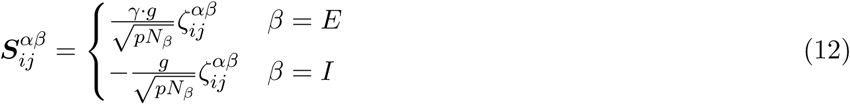

With *N_E_* excitatory neurons and *N_I_* inhibitory neurons, each neuron receives precisely *C_E_* = *pN_E_* excitatory connections and *C_I_* = *pN_I_* inhibitory connections from the rest of the network where *p* is the degree of sparsity in the network. The variable 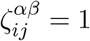 if a connection is present between neuron *j* in population *β* (presynaptic) and neuron *i* in population *α* (postsynaptic), and 0 otherwise. The variable *g* controls the coupling strength while 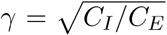 is used to balance the weight matrix in cases where *C_I_* ≠ *C_E_*. Here however, we do not train a balanced E/I network, but rather a balanced I network (see Materials and Methods: FORCE Training a Balanced I Network). We include this section for completion.

In normal FORCE training, the learned term is given by the following:

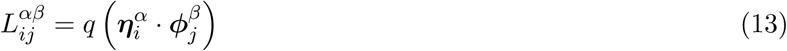

where *q* determines the amount of learned recurrence the network receives and 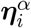 and 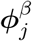 are referred to as the neural encoders and decoders, respectively [Nicola and Clopath, 2017]. The encoders help determine the tuning preferences of the neurons. The decoders are trained using Recursive Least Squares (RLS), an online *L*_2_ minimization scheme described in further detail in the section Recursive Least Squares. The network approximant of the intended dynamics is given by the following

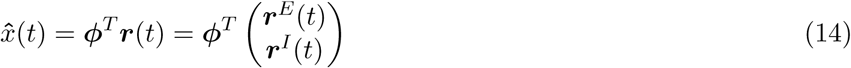

However, the weight matrix generated by this approach does not necessarily satisfy Dale’s Law, the constraint that a neuron cannot make both excitatory and inhibitory connections. While the static component of the weight matrix satisfies Dale’s law, the sum of the static and learned components do not necessarily satisfy this constraint. It is sufficient (but not necessary) to constrain the learned component of the weight matrix to independently satisfy Dale’s Law:

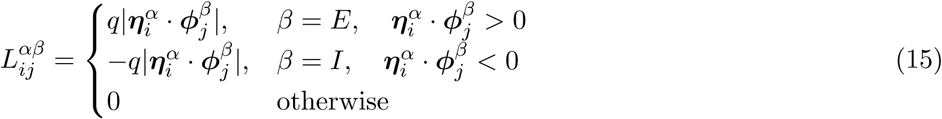

While implementing the boundary condition (15) on 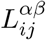 is sufficient to satisfy Dale’s Law, it is time consuming during learning as it requires *O*(*N*^2^) computations to bound every weight in the weight matrix. We opt to impose a condition on the encoders and decoders separately to enforce Dale’s Law. Consider the following operations:

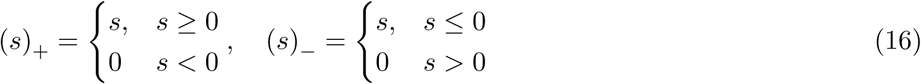

where the operation (*s*)_*±*_ sets *s* to 0 if its negative (positive) and retains its value otherwise.

First, we decompose the decoders and encoders as follows:

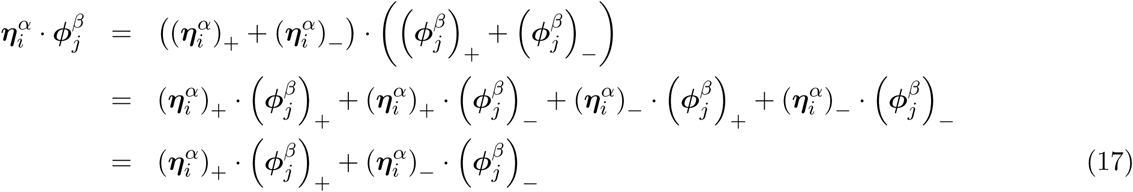

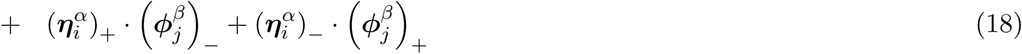

Note that the terms on line (17) are positive while the terms on line (18) are negative. Thus, we can consider the following weights

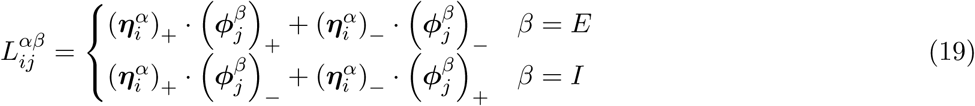

which implements an alternate boundary condition to (15) yet yields weight matrices that satisfy Dale’s Law. Furthermore, implementing (19) only requires *O*(*N*) computations as only encoders and decoders are bounded by their signs. This derivation, among other results, is originally owed to [Sauvage, 2016].

### 4.3 Recursive Least Squares

The decoders are determined dynamically to minimize the squared error between the approximant and intended dynamics, ***e***(*t*) = 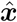(*t*) − ***x***(*t*). The Recursive Least Squares (RLS) technique updates the decoders accordingly:

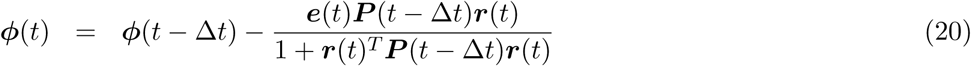

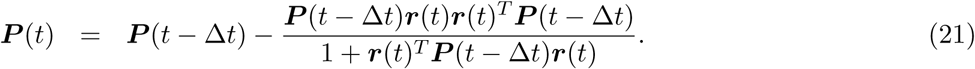

RLS is described in greater detail in [Sussillo and Abbott, 2009, Nicola and Clopath, 2017]. The network is initialized with *ϕ*(0) = **0**, ***P*** (0) = *I_N_ λ^−^*^1^, where *I_N_* is an *N*-dimensional identity matrix.

### 4.4 FORCE Training a Balanced I Network

In addition to the procedure described above, a balanced network can be generated with recurrent inhibition alone [Harish and Hansel, 2015]. In particular, we can consider the same network equations as before with the constraints:

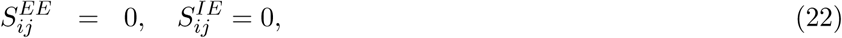

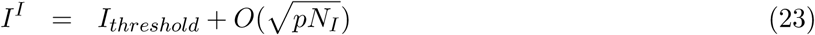

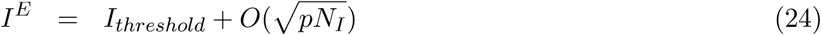

where the inhibitory network receives a super-threshold *I^I^* > *I_threshold_* = −40 pA background current that is being balanced by the recurrent inhibition. The excitatory neurons in the population also receive inhibitory connections and thus have a super-threshold background current after FORCE training is concluded. However, during FORCE training, the current *I^E^* is clamped to below threshold (*I^E^* < *I_threshold_*) to prevent firing in the excitatory neurons thereby eliminating both *EE* and *IE* weights. This occurs through the dependence of (20) on ***r***_*E*_ (*t*), and *ϕ*(0) = 0.

### 4.5 Internally Generated Theta Sequence

To construct a network with a recurrently generated theta sequence of firing, we FORCE train the network to learn the following supervisor:

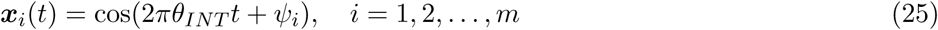

Here, *θ* is the frequency of the oscillation that is embedded in the recurrent weights of the network and *ψ_i_* is a phase shift for the *m* components of the supervisor. The *m* components of the supervisor yield an *N* × *m* dimensional matrix of encoders and decoders where *m* = 100 components. For each neuron, exactly one encoder element is non-zero and is set to 1. The phase shift is randomly selected and uniformly distributed over [0, 2*π*]. When trained in this fashion, the inhibitory currents and the septal inhibition combine to generate an interference pattern in the excitatory neurons (See Supplementary Figure S1):

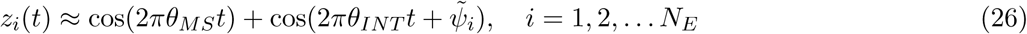

We interpret *z_i_*(*t*) to be the fluctuating component of the current an excitatory SHOT neuron receives. Note that we have written *ψ̃_i_* as opposed to *ψ_i_* as the phases of the excitatory neurons are only indirectly related to the phases in the supervisor (25). This is due to the operations applied in Equation (19).

### 4.6 Interference Based Control of the Population Activity by Septal Inputs

With the currents arriving at each neuron given by Equation (26), we consider the conditions under which the medial septum influences the population activity for the SHOT network. In particular, we will set out to provide sufficient conditions under which the population activity *ρ*(*t*) is a periodic function in time with period 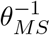. Proceeding generally, we assume that a network transforms its currents via some unspecified differentiable non-linearity *F* (*z*) where *F* (*z*) ≥ 0 into spike rates. This implies that the population activity is given by the following:

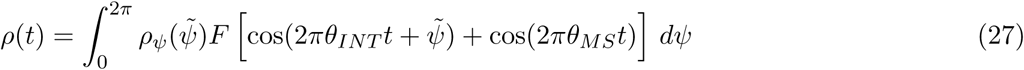

The function *ρ*(*t*) is the population activity for a network of neurons with identical tuning curves *F* (*z*) receiving heterogeneous currents, *z_i_*(*t*). The heterogeneity is in the phases of the currents, *z_i_*(*t*) through the variable *ψ̃_i_*. We assume that this parameter comes from a static distribution with density function *ρ_ψ_*(*ψ*). Now, let us consider 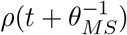:

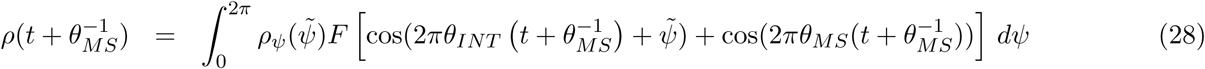

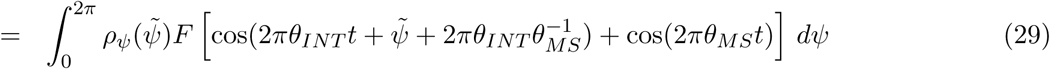

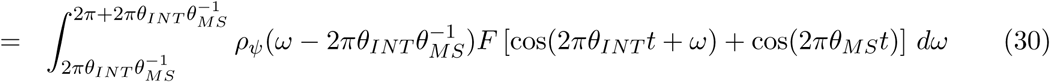

If the phase distribution is uniform (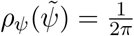), then:

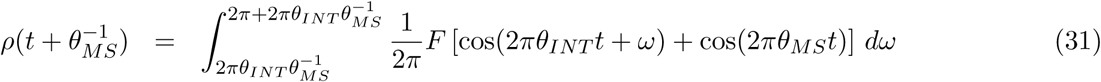

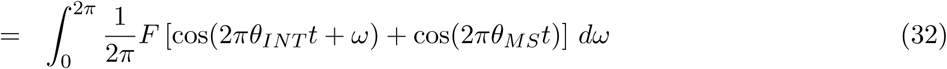

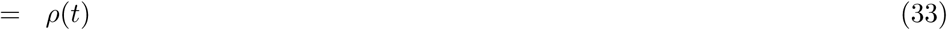

where line (32) is justified by the fact that *G*(*ω, t*) = *F* [cos(2*πθ_INT_ t* + *ω*) + cos(2*πθ_MS_t*)] is periodic in *ω* with period 2*π* and phase shifts (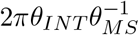 in line (31)) do not alter integrals of periodic functions. This implies that a uniform distribution is a sufficient condition for septal control of population activities. Finally, we consider what happens to *ρ*(*t*) when the septal inputs are removed:

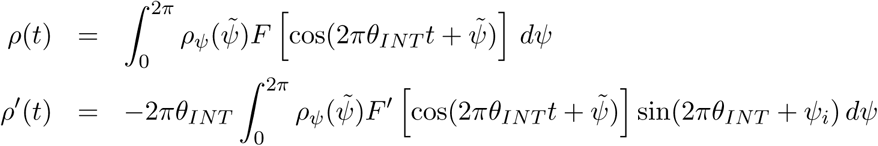

If we again assume that *ψ̃* is uniformly distributed, then we arrive at the following:

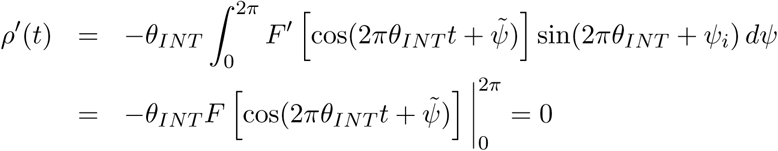

and thus *ρ*(*t*) is a constant and given by 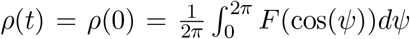. Thus for uniformly distributed phase preferences, the population activity is constant in the absence of septal inhibition despite the fact that all the neurons are oscillating.

### 4.7 Interference Based SPW-R Compression of Spike Sequences

The application of interference theory to explain phase precession is well studied [Burgess et al., 2007, O’keefe and Burgess, 2005, O’Keefe and Recce, 1993]. However, here we demonstrate that reversible sharp-wave ripple compression is also well explained by interference theory. Considering *z_i_*(*t*) again,

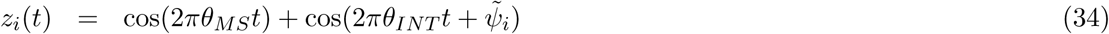

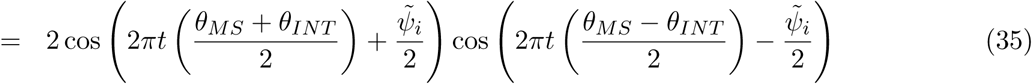

Equation (35) implies that the time-field order is controlled by the envelope 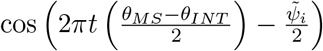 and in particular the phases *ψ̃_i_*. If the phase distribution of *ψ̃_i_* is uniform, then the firing-fields are uniformly distributed in time. If *θ_MS_* < *θ_INT_*, then the envelopes have phases *ψ̃_i_*/2 while if *θ_MS_* > *θ_INT_*, the phases become reversed with −*ψ̃*/2. The end effect of this transformation is to either preserve time-field order or reverse time field order. If septal inputs are removed, the interference pattern collapses and the neurons receive the input

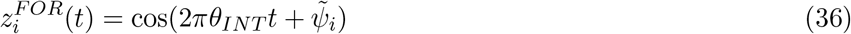

where *FOR* is short for forward compression. Removal of the septal inputs triggers a compression provided that the neurons also receive an additional super-threshold constant current. Thus, the default compression mode is a forward compression where the bursts of activity occur in the same order as the firing-fields when septal inputs are present. This compression can be reversed in two ways: 1) reverse the order of the firing fields *z_i_*(*t*) and maintain 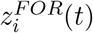 or 2) reverse 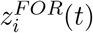 and maintain the order of firing fields.

In mirror reversion, the septal inputs are present and satisfy the frequency relationship *θ_MS_* > *θ_INT_*. The time fields are reversed in order relative to the compression phase. However, this is not a valid mechanism for reverse compression as if *θ_MS_* > *θ_INT_*, both excitatory and inhibitory neuron populations regress in phase. To our knowledge, phase regression has not been previously observed experimentally. In harmonic reversion, the firing-fields reverse near the harmonics of *θ_INT_* (see Supplementary Figure S3A). For *θ_MS_* just slightly larger than *θ_INT_* /2, the firing fields reverse and display phase precession instead of phase regression.

Finally, directly reversing 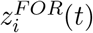 also triggers a compressed, reversed replay. An additional population of interneurons is sufficient for this type of reversion and thus we call it Interneuron Reversion. Reverse compression corresponds to the excitatory neurons in the SHOT network receiving the current:

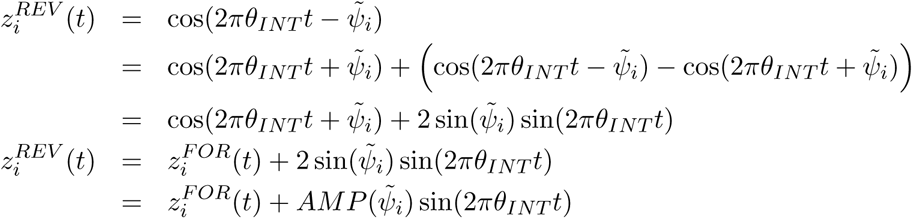

Thus, the reversion interneurons should be inhibited by the SHOT interneurons (through sin(2*πθ_INT_ t*)) while subsequently inhibiting the excitatory neurons. The amount of inhibition an excitatory neuron in the SHOT network receives from the reversion population is controlled by the amplitude function *AMP* (*ψ̃_i_*) = 2 sin(*ψ̃_i_*).

### 4.8 Reversion Interneuron Population

The reversion population is a population of *N_REV_* interneurons with LIF dynamics that receives inhibition from SHOT interneurons. For simplicity, we take *N_REV_* = *N_I_* and 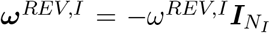, where 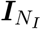 is the *N_I_* × *N_I_* identity matrix and *ω^REV,I^* is a scalar quantity with (*ω^REV,I^* = 1) that determines the amount of inhibition the reversion population receives. The only parameter difference between the reversion population and the other neurons is the background current, *I^REV^* which varies when the GABAergic septal current is present or absent (see Figure 2, Figure 4 Captions).

From the preceding section, we require the reversion population to transmit the signal 2 sin(*ψ̃i*) sin(2*πθ_INT_ t*) in a biologically plausible manner. First, the reversion interneurons must be locked into the internally generated theta oscillation through the term sin(2*πθ_INT_ t*). Thus, we train a decoder *ϕ^REV^* that can decode sin(2*πθ_INT_ t*) from the activity of the reversion population:

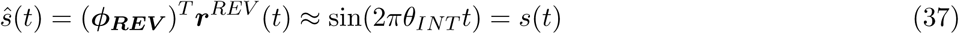

where *ϕ_**REV**_* is trained using the decoded internal oscillators of the SHOT network as a supervisor. The training is not performed online as in RLS but is learned immediately through *L*_2_ optimization:

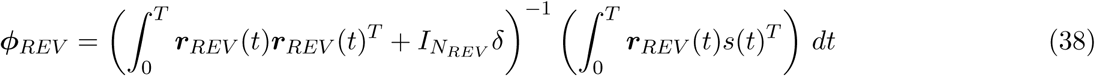

where *δ* = 50 acts as a regularization parameter and *T* = 25 s was used to compute *ϕ^REV^*. Finally note that the current:

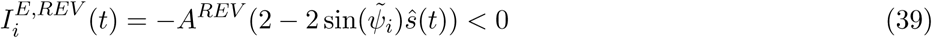

is strictly inhibitory and performs the necessary reversion in the excitatory neurons. The parameter *A^REV^* is used to scale the current upward so that the SHOT interneuron and reversion interneuron currents are of similar magnitude (*A^REV^* = 10 pA). As the excitatory neurons receive an extra source of inhibition, they require a larger excitatory drive when the septal inputs are deactivated. Further, as the current 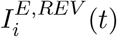 is strictly negative, a set of weights ***ω***^*E,REV*^ can be found by a constrained optimization problem

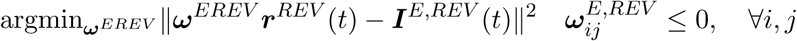

This problem can be solved numerically for example with the MATLAB function lsqnonneg. However, for simplicity, we do not take this approach and simply apply the negative current (39) to the excitatory SHOT neurons.

### 4.9 Gamma-Assembly Frequency as a Function of *θ_INT_* and *θ_MS_*

Here, we will derive the relationship between the septal frequency, *θ_MS_*, the interneuron recurrent oscillation frequency, *θ_INT_*, and the gamma-assembly frequency during compressed replay (without septal inhibition) which we refer to as Γ. The quantity *n_x_* denotes the number of cycles of the oscilltion *x* (either *x* = *MS* or *x* = *INT*) in a specific time period while 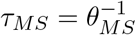, 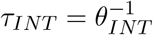, and *τ* = (*θ_INT_* − *θ_MS_*)^−1^ is the period of the time-field firing. First, as the number of theta cycles (*θ_MS_*), determines the number of gamma assemblies (Figure 5F), then the total number of theta cycles is

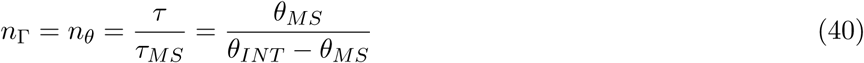

then the frequency of gamma during SPW-R compression, which we define as 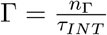 is given by

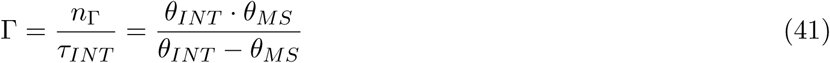

As our analysis in the Methods section Interference Based Control of the Population Activity by Septal Inputs demonstrates, we only observe *θ_MS_* in the population activity or other aggregate/macroscopic metrics. Thus, it is useful to write *θ_INT_* = *θ_MS_* + ϵ and thus:

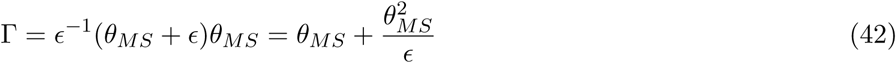

which in our case yields ϵ = 0.5 Hz, *θ_MS_* = 8 Hz, yields a gamma assembly frequency of Γ = 136 Hz. This is in good agreement with the results of Figure 4 and Figure 5. In Figure 4, we see 13 assemblies over an 88 ms interval, yielding Γ ≈ 147.7 Hz (forward replay), 13 assemblies over 98 ms, yielding Γ ≈ 132.65 (reverse replay), while in Figure 5 we see 8 assemblies over a 70 ms interval yielding Γ ≈ 114 Hz.

The gamma frequency Γ corresponds to the number of assemblies per unit time in the recurrent *θ_INT_* oscillation. During theta-nested on gamma situations when septal inputs are on, the number of gamma-assemblies will always be less than (or equal to) the number observed during a ripple. Further, a neuron in the network spikes with a frequency of 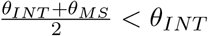. Thus, the gamma-assemblies that do occur are still dilated when septal inputs are present relative to when they are absent. As such, less assemblies occur and thus the gamma oscillation frequency, defined as the number of assemblies per unit time, is always faster for a ripple than during normal theta activity.

### Detailed Methods for Each Figure

#### Methods for Figure 1: FORCE Training a Septal Hippocampal Oscillatory Theta (SHOT)

A Leaky-Integrate-and-Fire network with an even split of excitatory and inhibitory neurons (see Table 1 for further details) is FORCE trained to learn an internal theta (with frequency *θ_INT_*) oscillation. The training occurs simultaneously to the presentation of an external oscillation (with frequency *θ_MS_*). However, the network is trained only once with an external input with *θ_MS_* = 8 Hz thereby creating 2 second long interference patterns. The internal frequency oscillation is set at *θ_INT_* = 8.5 Hz while the external frequency varies across figures. The interference pattern is caused by the recurrent inhibition and the external septal inhibition oscillating at slightly different theta frequencies.

#### Methods for Figure 2: Two Mechanisms of Reverse Compression

For Mirror Reversion, all parameters are identical to the normal compression case with the exception of *θ_MS_*, the septal oscillation. Here, the septal oscillation frequency is taken to by 9 Hz instead of the default 8 Hz used in the majority of simulations. The reversion interneurons are brought online simultaneous to removal of septal inhibition by applying an extra 30 pA current to the nominal value in Table 1. The reversion neurons receive inhibition from the inhibitory neurons in the SHOT network through the weight matrix ***ω***^*REV,I*^. Each reversion interneuron receives coupling from a single interneuron from the SHOT network. As the SHOT interneurons are not sorted with respect to their indices, we take 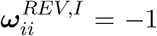 and 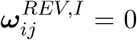, *i* ≠ *j*. The background current when they are not applied is set to the threshold value (−40 pA). Finally, the excitatory SHOT neurons receive an extra current valued at 40 pA (instead of 20 pA, as in Figure 1) due to the extra inhibition they receive from the reversion interneurons in this mechanism for reverse replay.

#### Methods for Figure 3: Learning Online with a Single Stimulus Presentation

Here, we decode the excitatory and inhibitory neurons with a time constant of *τ_D_* = 10 ms and *τ_R_* = 2 ms. As the spiking in the network is sparse and segregated to theta locked firing, to replay a continuous trajectory we introduced heterogeneity into the background firing rates for the neurons. Thus, we created copies of the inhibitory to excitatory weight matrix to increase the network size and add sparsity. The network consisted of 2000 inhibitory neurons and 12,000 excitatory neurons. The excitatory neurons were given fixed currents distributed over [−35, 5] pA with a uniform distribution. Compression was triggered by removal of the septal inputs and application of an additional 20 pA current to all neurons. Furthermore, the due to size constrains, the Fourier rule was not explicitly implemented. Instead, the *L*_2_ problem was solved offline using principal conjugate gradient descent (pcg function in MATLAB) with a regularization set at *λ* = 10^5^ using the time interval [4, 6] to generate the basis ***r***(*t*). However, a subset of the weights were computed online to demonstrate how the rule can also be implemented online.

#### Methods for Figure 4: Rapid Spike Train Learning with a Septal-Hippocampal Oscillatory Theta Network

A population of *N_RO_* = 2000 Read Out (RO) neurons were stimulated with a sequential current that activated each neuron for 40 ms with a pulse height of 10 pA:

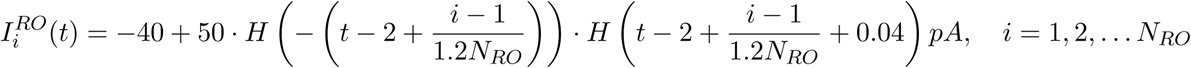

where *H*(*t*) is the Heaviside function. Note that the static component (−40 pA) is the baseline current the RO neurons always receive (and is not in addition to the amount in Table 1). To generate the external inputs in Figure 4F, we applied the same current as in

The *λ* parameter for the local Fourier Rule was *λ* = 4.25 · 10^−7^. Unlike in Figure 3, the Fourier rule is run online here only without delay term implemented. If the spike-train supervisor is less than the time periodicity (2 seconds), than the forgetting term is unnecessary. The excitatory neurons received an additional current of 18 pA when septal inputs were removed during forward replay and 40 pA during reverse replay.

#### Methods for Figure 5: Rapid Spike Train Learning with CA1 Constrained Connectivity

The inhibitory neurons in CA1 received strong all-to-all connectivity from the excitatory population with 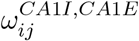 was randomly drawn from a uniform distribution on [0, 25/*N*_*CA*1*E*_]. The excitatory neurons likewise received all-to-all connectivity from the inhibitory population with 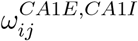 drawn from a uniform distribution on [−0.03/*N*_*CA*1*I*_, 0]. This however was not necessary for the results of Figure 5. The local Fourier rule rate parameter was *λ* = 7.875 · 10^−7^. The input from the entorhinal cortex follows from Figure 4:

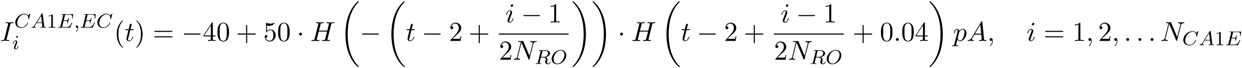

The excitatory neurons in the CA3 network receive an extra 22 pA current when the septal inputs are off.

For Figure 5J, the excitatory neurons in the CA3 network receive an extra 20 pA applied current as a stochastic process. At each time step, *dt* the ripple inducing current transitions from 0 pA to 20 pA with a constant probability *p* = *p*_0_ · *dt* = 0.00025 where *p*_0_ = 5 Hz sets the background ripple rate. Without any inhibitory modulation, this corresponds to a poisson process for generating SPW-Rs. If the current transitions from 0 to 20 pA at time *t*\*, it stays in that state until 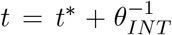. After this cycle, the extra current transitions back to 0 pA. A relative refractory period is applied for the subsequent theta cycle where the probability of initiating another SPW-R varies is decreased:

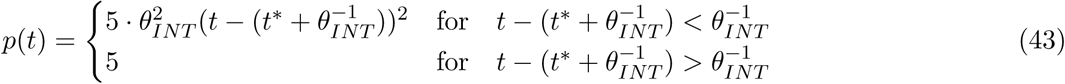

Note that the term 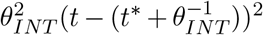 scales between 0 and 1 on 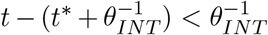. Additionally, the incorporation of the relative refractory period only alters the inter-ripple-interval distribution on [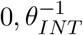] but will not induce harmonics. For Figure 5K, the same approach is applied only the probability of transition *p*(*t*) now always varies in time instead of just at relative refractory periods. We take the probability to be theta-modulated and dependent on the CA3 interneuron population at each time step, *dt*: *p*(*t*)·*dt* = (5.1+2.5·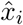(*t*))·*dt* where 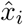(*t*) is one of the FORCE trained decoded oscillators from the inhibitory neurons in CA3. This model has two parameters, a constant baseline probability and the amplitude of the oscillatory components. For convenience, we took the first element, *i* = 1 and ignored effects on the fragmentation error as for this figure, we are only assessing inter-ripple-interval distributions. The refractory period is applied as before:

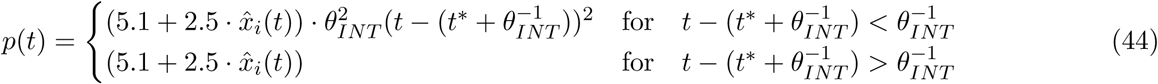

The network is allowed to run for 3000 seconds of simulation time. The inter-ripple-interval distribution was computed as a kernel density estimator over [0, 1] s with a bandwidth of 10 ms using the MATLAB function ksdensity. Finally, the inter-ripple-interval was measured using the discrete times 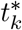, with 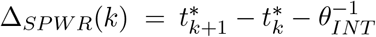 where 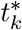 is the *k*th time the ripple is induced by the extra current.

## Supplementary Material

**Supplementary Figure S1:**
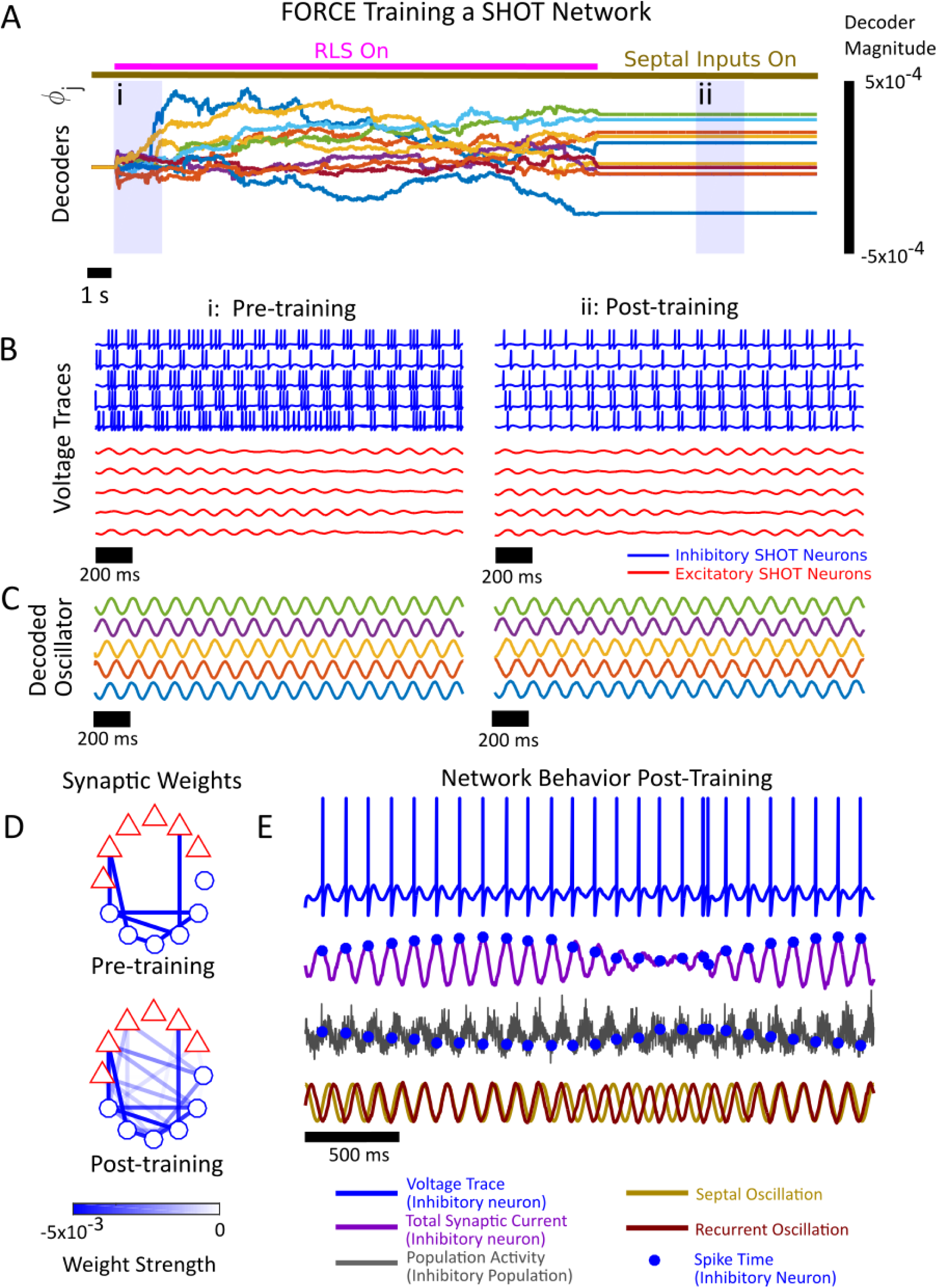
FORCE Training the Septal Hippocampal Oscillatory Theta (SHOT) Network. A network consisting of 2000 Excitatory and Inhibitory LIF neurons was FORCE trained to oscillate at its own internal frequency, *θ_INT_*. **(A)** A series of decoders is learned dynamically through Recursive Least Squares (RLS) minimization. The decoders are used to decode the signal cos(2*πθ_INT_ t* + *ψ_i_*) for *ψ_i_* drawn from a uniform distribution on [0, 2*π*] for *i* = 1, 2, … 100 components. The network receives the inputs from the medial septum while it is trained. The decoders also function to stabilize the network dynamics through a learned feedback weight (see Materials and Methods for further details). The overlines denote the presence of septal inputs (gold) or RLS learning (pink). The evolution of 10 decoders is shown. **(B)** Voltage traces for 5 excitatory neurons (red) and 5 inhibitory neurons (blue) are shown for the epochs (i)-(ii) from (A). The excitatory neurons were kept sub-threshold by applying a low background current which was insufficient to induce spiking (*I^E^* = −25*pA*). The inhibitory neurons fire either bursts or single action potentials during a theta oscillation while the excitatory neurons display a subthreshold interference pattern before and after training when MS inputs are present. **(C)** The intrinsic oscillator cos(2*πθ_INT_ t* + *ψ_i_*) can be decoded during training (i, bottom) and after training (ii, bottom) from the filtered spike times, ***r***(*t*), of the inhibitory neurons. A subset of 5 decoded oscillators are shown. **(D)** A subset of 5 excitatory (red) and 5 inhibitory (blue) neurons prior to FORCE training (top) and after FORCE training (bottom). No excitatory connection is present in this network either before or after training. The learned component of the inhibitory coupling is a perturbation to the initial random coupling. **(E)** Voltage trace (blue) for an inhibitory neuron along with the total current the neuron receives (purple). The inhibitory population activity (grey) shows a strong theta modulation that is unaffected by the interference pattern from the septal and recurrent inhibitory oscillations (gold and red, respectively). The spike times for the inhibitory neurons (blue dots) are locked to the total synaptic current and precess in phase with respect to the population activity.

**Supplementary Figure S2:**
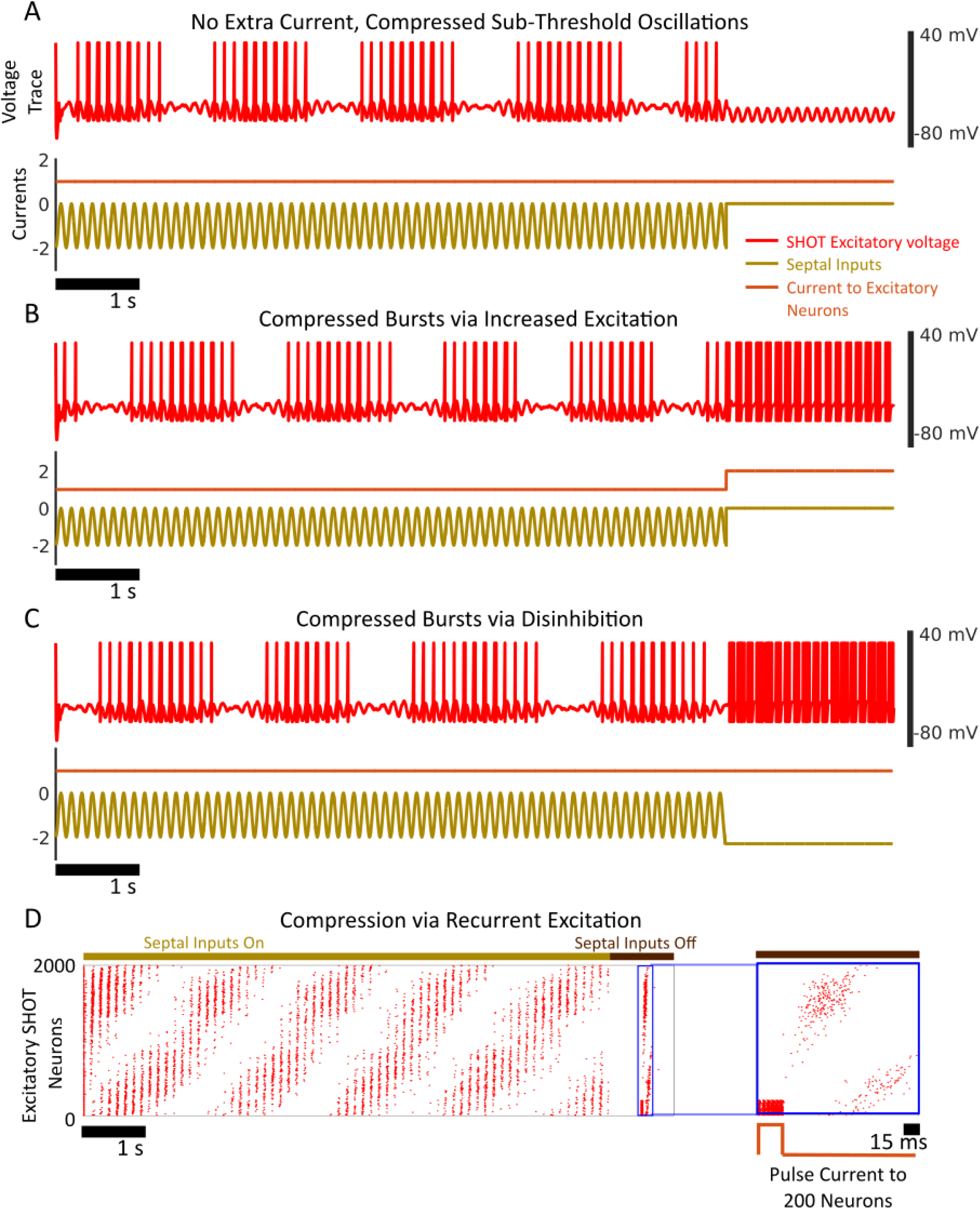
Removing Septal Inputs and Triggering Compressed Bursting. **(A)** (Top) Voltage trace for an excitatory neuron (red) in the Septal Hippocampal Oscillatory Theta (SHOT) network, (Bottom) the background current on the excitatory neurons (orange) and the septal inhibition (gold) arriving at the inhibitory neurons. If the septal inputs are turned off (during the last 2 seconds), subthreshold oscillations emerge due to the recurrent inhibition in the Septal-Hippocampal Oscillatory Theta (SHOT) network. The excitatory SHOT neurons only receive a constant background current (*I^E^* = −5 pA). **(B)** The subthreshold oscillations can be made superthreshold by providing the excitatory neurons an extra excitatory current (valued at 20 pA during last two seconds, bottom). **(C)** Alternatively, the excitatory neurons can also be disinhibited to trigger compressed bursts. In this implementation, the inhibitory neurons receive more septal inhibition in the last two seconds. The septal inhibition in the last two seconds however is static, and non-oscillatory and valued at −22 pA. Note that the septal and excitatory currents are not to scale here. **(D)** Recurrent connections between the excitatory SHOT neurons can also trigger compression. Here, the excitatory neurons (spike times in red dots) were coupled in an all-to-all manner with 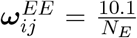, for all *i*, *j*, 1, 2, …, *N_E_* while they also projected to the inhibitory neurons in an all-to-all manner, 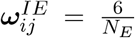 for *i* = 1, 2, … *N_I_* and *j* = 1, 2, … *N_E_*. The connections here were implemented with a long synaptic filter, *τ_D_* = 100 ms and *τ_R_* = 10 ms, similar to the time constants of NMDA synapses. A subset of 200 neurons was stimulated with a short pulse current of width 25 ms and peak 145 pA while the septal inputs were off (brown overline). To elicit spiking, the excitatory neurons received a white noise current with a standard deviation of 8 pA. This was enough to elicit spiking when septal inputs were on (gold overline, first 8 seconds) but not when septal inputs were off (last 1 second). Furthermore, the noise induced (rather than mean-driven) firing prevented excitatory feedback loops amongst the excitatory SHOT neurons due to the strong recurrence. The pulse current to the subset of excitatory SHOT neurons in conjunction with recurrent excitation is enough to trigger a compressed replay (right, zoomed in segment).

**Supplementary Figure S3:**
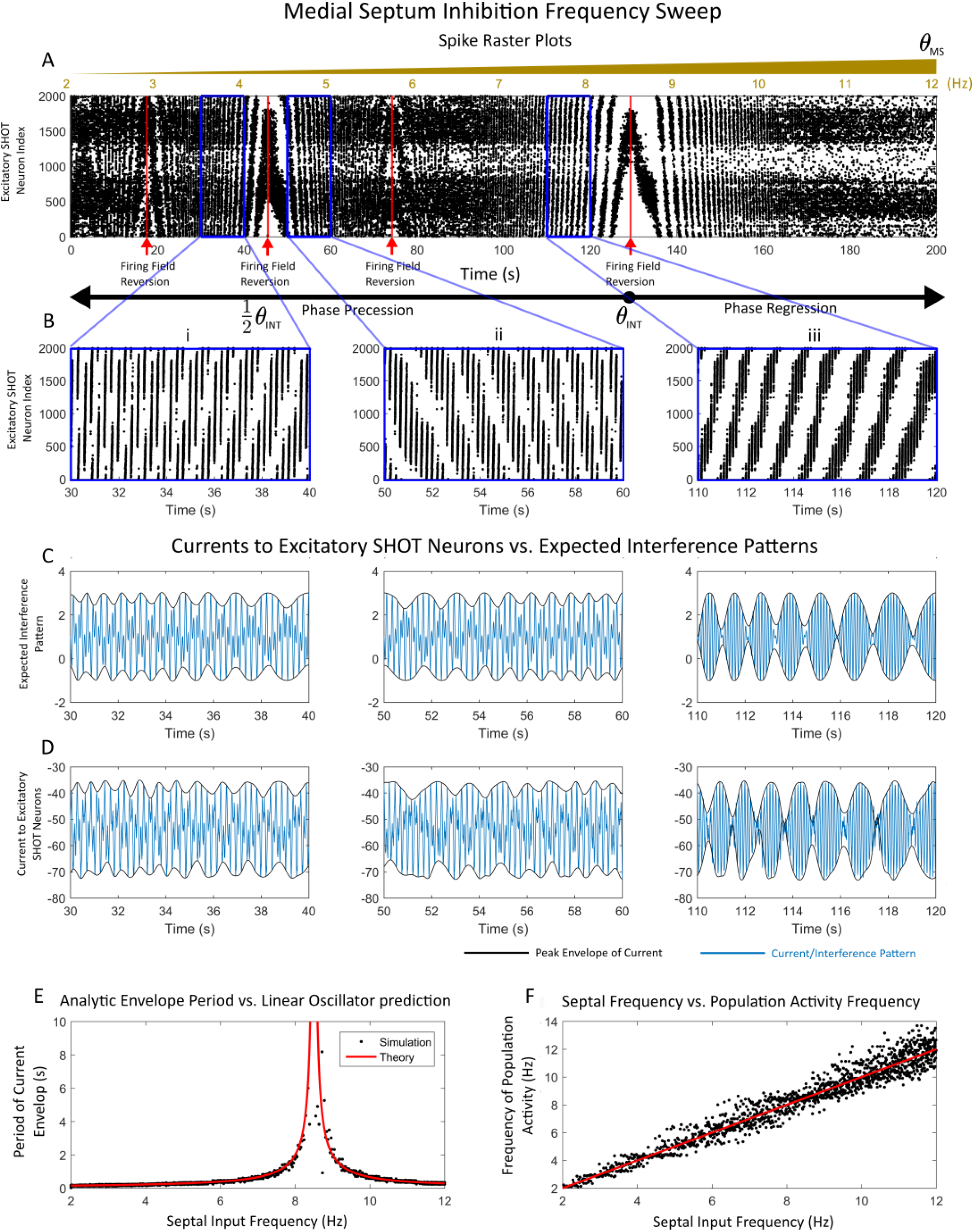
Determining Robustness of the SHOT Network to Septal Frequency Modulation. The frequency of the Medial Septum (MS) GABAergic inputs is swept from 2 Hz to 12 Hz over a 200 second interval using the MATLAB chirp function. **(A)** The spike raster plots of the excitatory SHOT neurons are sorted according to increasing phase preference. Note that the bottom axis corresponds to time while the top axis corresponds to frequency of the MS inputs. The gold overline denotes the presence of septal inhibition. The three segments (i)(30, 40)*s* (ii)(50, 60)*s* and (iii) (110, 120)*s* are highlighted for subsequent analysis in (B)-(D). The time-fields periodically flip in orientation near harmonics of the recurrent frequency *θ_INT_* = 8.5*Hz*. Note that time cells fire with slightly different rates within a firing field. This is likely due to errors in FORCE training. **(B)** The zoomed raster plots for the segments outlined in (i)-(iii). Note that the time-fields flip in orientation on either side of the harmonics. **(C)** The peak envelope (black) and the interference pattern (blue) generated by adding the septal input to a perfect linear oscillator with a frequency of *θ_INT_* for the segments (i)-(iii) outlined in (A). **(D)** The currents (blue) arriving at the excitatory neurons from the inhibitory interneurons in addition to the peak envelope (black). Note that the currents closely correspond to the expected behavior with perfect linear oscillators in (C). The envelope is computed using the MATLAB envelope function for (C)-(D) using the peakfinder function (see MATLAB file exchange). **(E)** The period of the envelope for the network (black) and the expected value (red) when the envelope is computed using the Hilbert transform. The envelope is computed for the input currents for one of the excitatory neurons. Peak to peak differences in the envelope are used to determine the periodicity of the envelope function. For MS frequencies near *θ_INT_*, this corresponds to the periodicity of the time-fields. A different formula is required near the harmonics. **(F)** The frequency of the population activity as a function of the input frequency of the MS. The frequency is estimated using the inhibitory interneurons as they have faster firing rates distributed over the entire-theta interval, thereby generating a smoother estimate. The peak to peak time interval is subsequently measured to compute the instantaneous frequency.

**Supplementary Figure S4:**
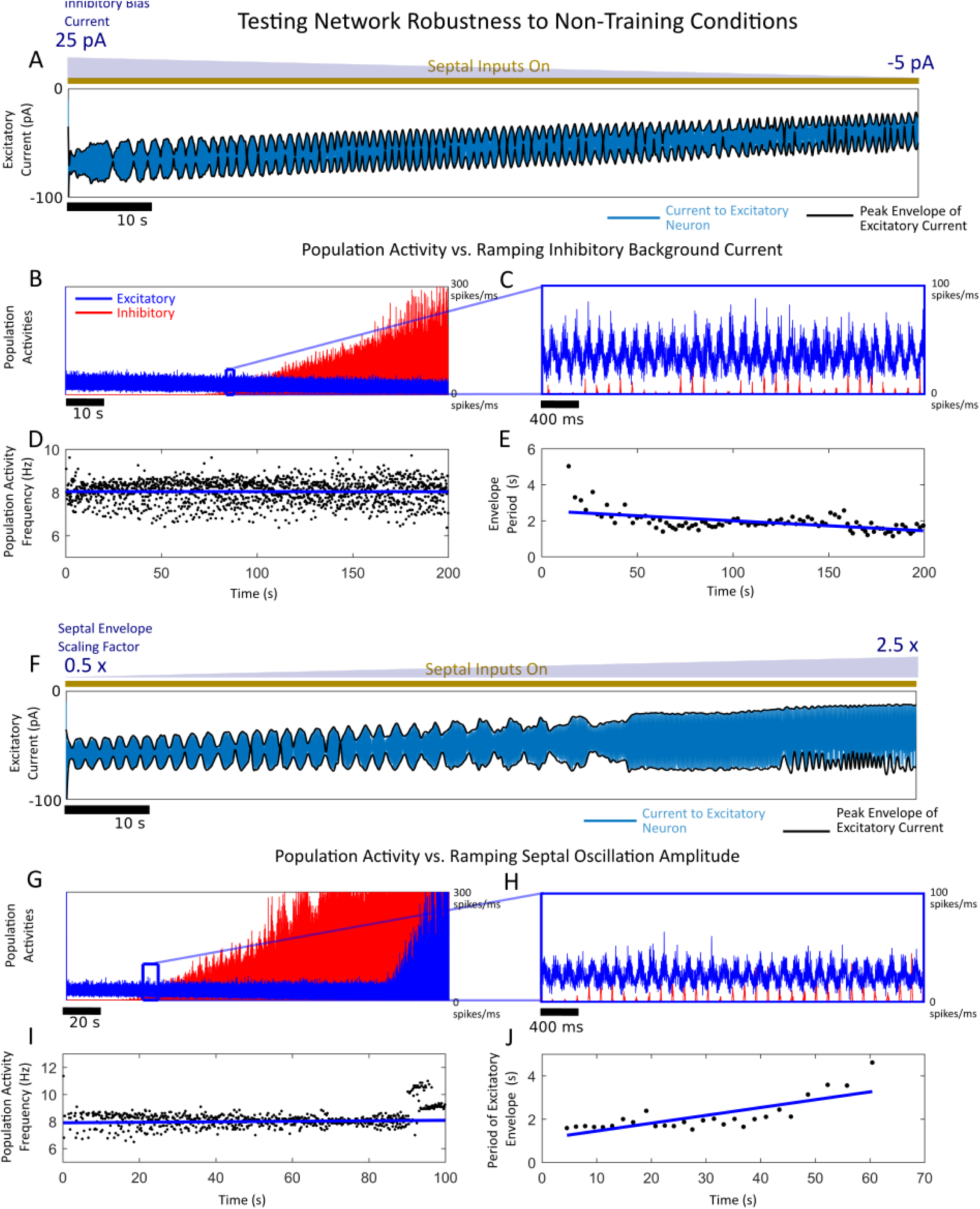
Determining Robustness of the SHOT Network to Septal Amplitude Modulation. For (A)-(E), the background current to the inhibitory SHOT neurons (*I^I^*) was linearly ramped relative to the baseline amount over a 200 second interval. This corresponds to a range of [−25, 5] pA of applied current. For (F)-(J), the Medial Septum GABAergic amplitude parameter (*I^GABA^*) was linearly ramped from 0.5× to 2.5× the baseline input amplitude (-5 pA to -25 pA) over a 100 second interval. (**A**) The current arriving at an excitatory SHOT neuron (teal) displays an interference pattern (envelope in black) for all current levels. Higher or lower levels of excitatory drive are required to induce spiking for a given level of applied current to the inhibitory neurons. (**B**) The population activity for the excitatory neurons (red) and inhibitory neurons (blue). As the background current increases, the excitatory neurons increase their firing through disinhibition. (**C**) A zoomed in, 2 second segment from (B). (**D**) The frequency of the theta oscillation was measured by measuring the inter-peak interval. Linear regression revealed a small negative slope (−3.182 · 10^−5^) that was not statistically significant (*p* = 0.89341). (**E**) Period of the interference pattern (black dot) vs the least squares fit (blue line). Linear regression revealed a decreasing trend in the interference pattern periodicity with a slope of (−0.00556) that was significant (*p* = 1.22 · 10^−9^). (**F**) The current arriving at an excitatory neuron (blue) displays an interference pattern (envelope in black) for moderate levels of MS GABAergic oscillation amplitude up to approximately 1.5×. (**G**) The population activity for the excitatory neurons (red) and inhibitory neurons (blue). As the GABAergic amplitude increases the excitatory neurons initially increase their firing through disinhibition. Past a critical point, the behavior of the SHOT network breaks down and the interneurons and excitatory neurons fire highly synchronized and pathological bursts. (**H**) A 2 second close up from (G). (**I**) The frequency of the theta oscillation in the population activity as a function of time. Linear regression revealed a small negative slope (−0.00186 · 10^−5^) that was statistically significant (*p* = 0.0186). Note that the pathological segments were removed (*t* > 80) in performing this analysis. (**J**) Period of the interference pattern (black dot) vs the least squares fit (blue line). Linear regression revealed an increasing trend in the interference pattern periodicity with a slope of (0.036184) that was significant (*p* = 2.38 · 10^−6^).

**Supplementary Figure S5:**
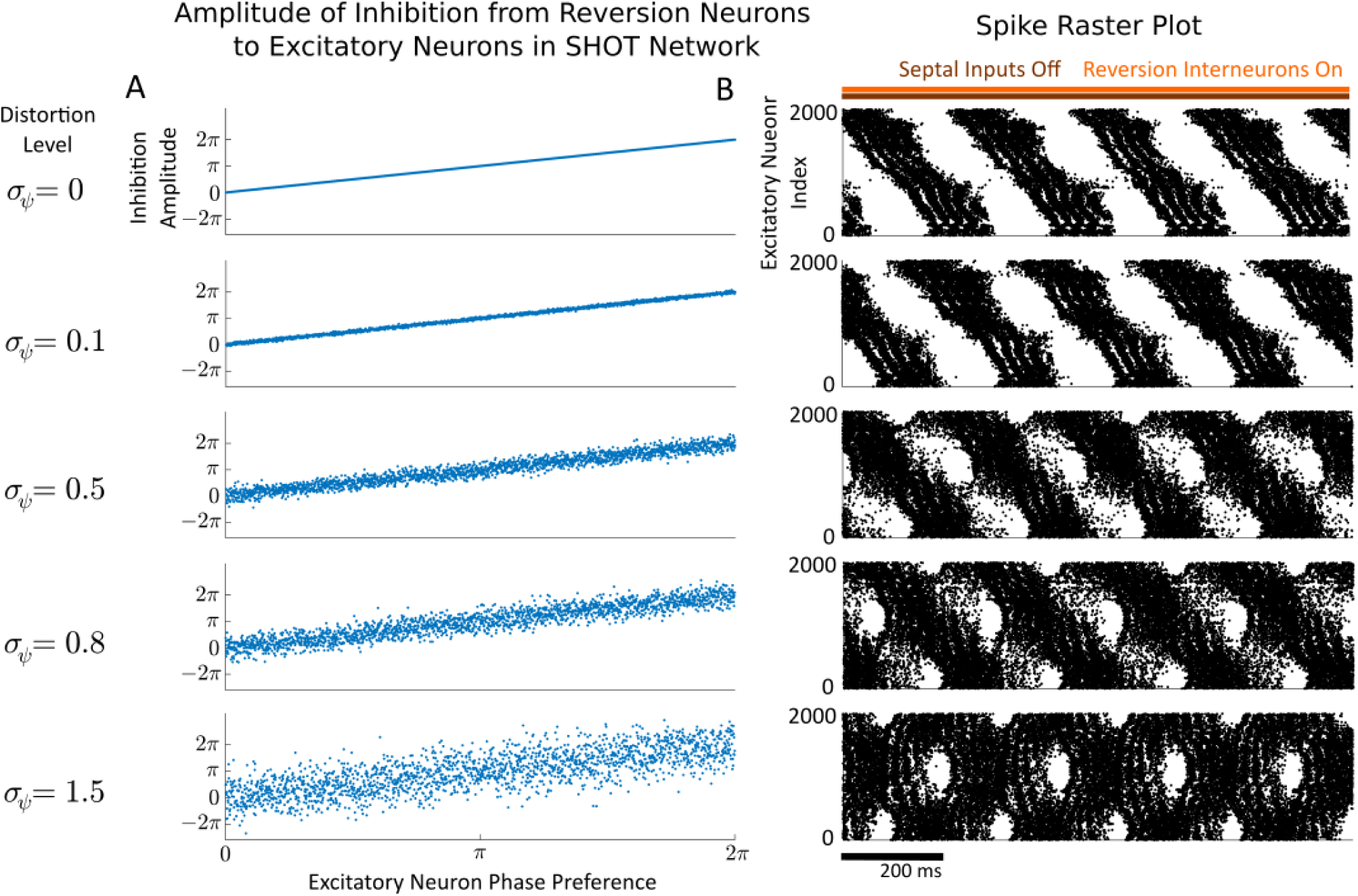
Determining Robustness of Interneuron Reversion to Amplitude Distortions. **(A)** Heterogeneity was added to the amplitude-phase relationship between inhibition from the reversion interneurons and the phase preference of the excitatory SHOT neurons. In particular, the functional relationship between the phase preference of the excitatory neurons (*ψ̃_i_*,x-axis) was corrupted by adding a 0-mean normally distributed noise term with standard deviation *σ_ψ_* to *ψ̃_i_* for 5 values *σ_ψ_* = 0, 0.1, 0, 5, 0.8, 1.5. **(B)** The raster plot for the excitatory neurons with septal-inputs off at the corresponding noise level on the left. The network was simulated for a 1 second transient before plotting. For increasing noise levels, the reversion becomes corrupted by a forward replay, the default firing of the network.

**Supplementary Figure S6:**
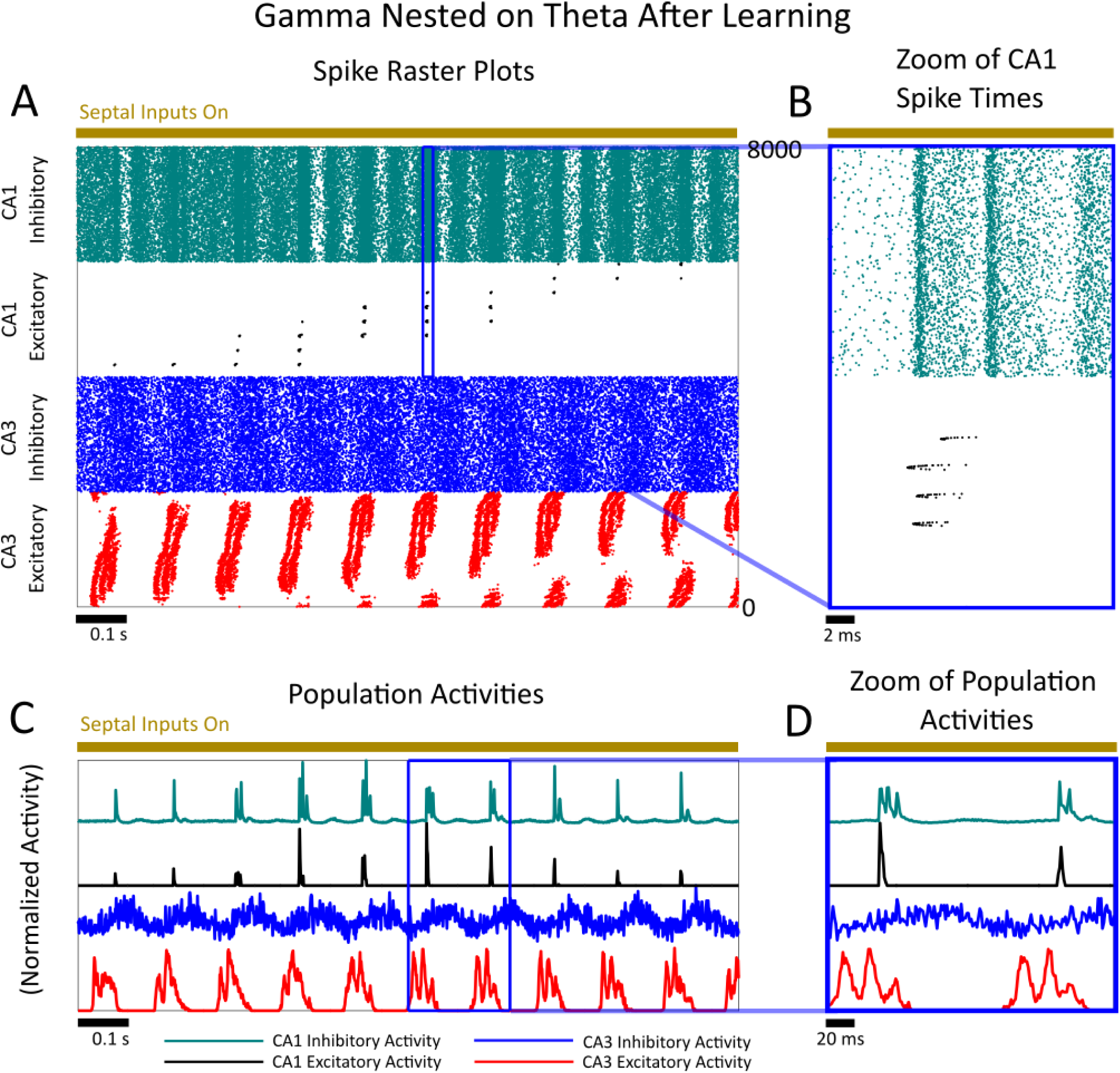
Gamma Nested on Theta Oscillations After Learning in the Presence of Septal Inhibition. **(A)** The CA3/CA1 network considered was trained identically as in Figure 5, with the local Fourier rule. The spike raster plot is shown for the CA1 inhibitory neurons (teal), the CA1 excitatory neurons (black), the CA3 inhibitory neurons (blue) and the CA3 excitatory neurons (red). Here, the septal inputs are kept on after learning and a small additional current (valued at 2 pA) is given to the excitatory CA3 neurons. This increases their activity rate, triggering gamma assemblies in CA1. These assemblies are nested on the theta oscillation. The assemblies repeat on the same time scale as they were originally observed (see Figure 5). **(B)** A zoomed view of the CA1 population spike times demonstrating gamma assemblies in the excitatory neurons and gamma frequency firing in the inhibitory neurons. **(C)** The scaled population activities for the CA1 inhibitory (teal), the CA1 excitatory (black), the CA3 inhibitory (blue) and the CA3 excitatory (red) populations. The population activity in all cases is computed as a histogram with 1 ms time bins. **(D)** A 200 ms zoom of the population activity from (C). The gamma frequency activity is nested to the theta oscillation.

### Supplementary Material Section S1: Derivation of the Local and Global Fourier Learning Rules

Here, we will demonstrate how a Hebbian learning rule can be sufficient for single-instance learning of spike trains and behavioral supervisors. One mechanism for learning immediately is to regress onto a pre-existing and repeating (oscillatory) spike train. First, if we consider the signal *x*(*t*) that our system should remember, it can be bound to an underlying oscillation ***r***(*t*) through a set of decoders or weights, *ϕ* on [0, *τ*] where *τ* is the period of the oscillation. This yields the following equation for the decoders:

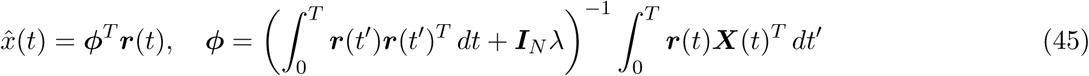

where ***X***(*t*) is the intended supervisor and ***r***(*t*) is the filtered spike train (9)-(10). The formula for the decoder is determined through the *L*_2_ on the following cost function:

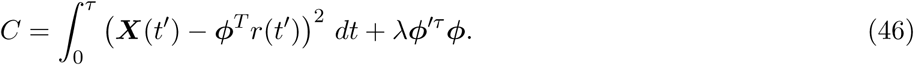

This approach is not plausible as it requires storage of ***X***(*t*) to compute *ϕ* through equation (45). Furthermore, this rule limits us to the interval [0, *τ*]. Thus, using Recursive Least Squares as inspiration, we rewrote this rule in an online fashion under the assumption that ***r***(*t*) is oscillatory. In particular, suppose that *ϕ* explicitly depends on time, *ϕ* (*t*) and resolves the following optimization problem:

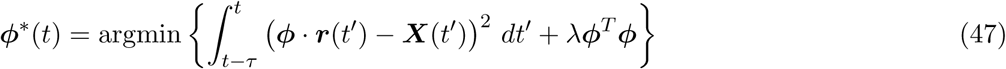

In other words, *ϕ*\*(*t*) give us the decomposition of *x*(*t*) on [*t* − *τ*, *t*] using the oscillatory backbone ***r***(*t*) as a temporal basis. If we solve the *L*_2_ problem, we immediately have the following:

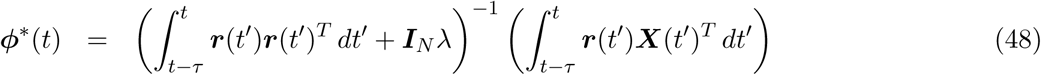

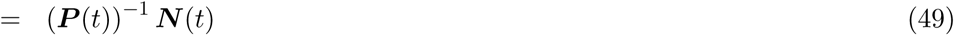

where

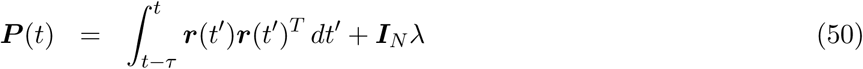

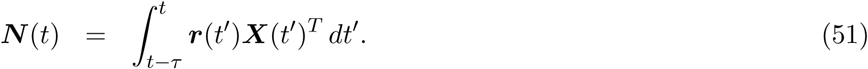

Now, we will consider the dynamics of ***P*** (*t*) and ***N*** (*t*) separately. First, the dynamics of ***P*** (*t*) become:

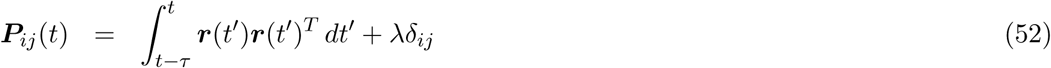

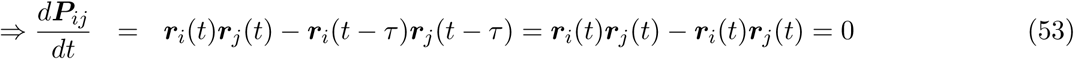

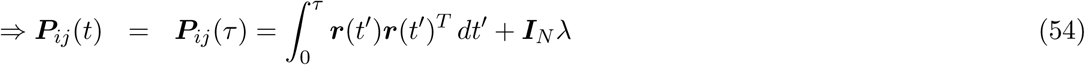

where *δ_ij_* is the Kronecker delta. This implies that ***P*** (*t*) is a constant for *t* > *τ*. The dynamics for ***N*** (*t*) become:

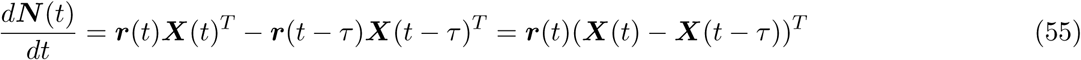

where we have used the oscillatory nature of ***r***(*t*) to simplify. This yields the online decoder dynamics:

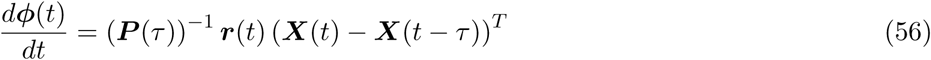

This yields the following learning rule which we refer to as the non-local Fourier rule.

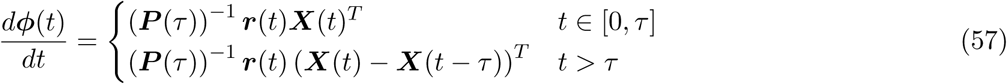

Let us reconsider the equation for the approximant, 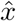(*t*)

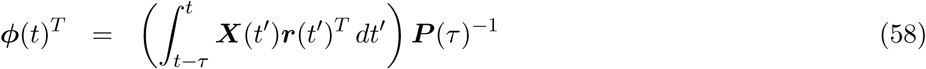

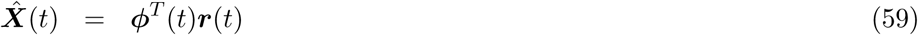

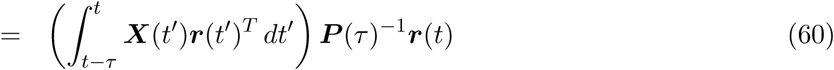

As the matrix ***P*** (*τ*) is symmetric and positive definite (***P*** (*τ*) is a Gramian matrix with a positive definite diagonal perturbation ***I***_*N*_ *λ*), it decomposes as ***P*** (*τ*) = ***U***^*T*^ ***DU*** where ***U***^−1^ = ***U***^*T*^ and ***D*** is a diagonal matrix with ***D***_*ii*_ > 0. Thus, the transform ***μ***(*t*) = ***Ur***(*t*) naturally maps our original basis into an orthogonal basis ***μ***(*t*). In particular, first note that if ***ν*** is an eigenvector of ***P*** (*τ*) with eigenvalue of *β* then ***ν*** is also an eigenvector of ***P*** (*τ*) + ***I***_*N*_ *λ* with eigenvalue *β* + *λ*. This immediately implies the following:

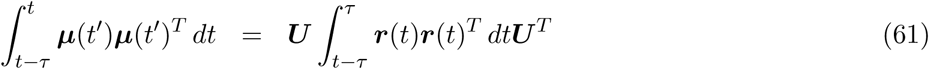

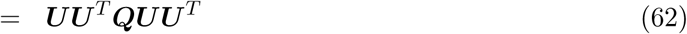

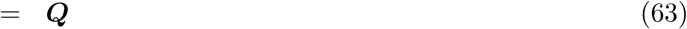

where ***Q*** is the diagonalization of the Gramian matrix, 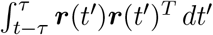 and ***Q*** + ***I***_*N*_ *λ* = ***D***. Thus, ***μ***(*t*) is orthogonal. This yields the following:

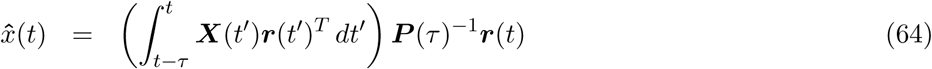

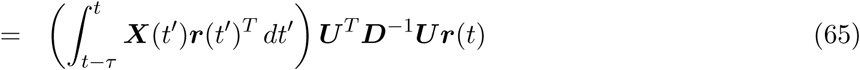

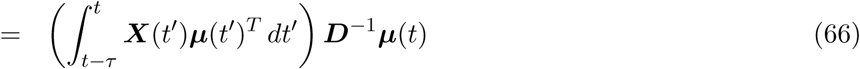

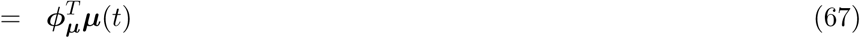

Note which yields the following:

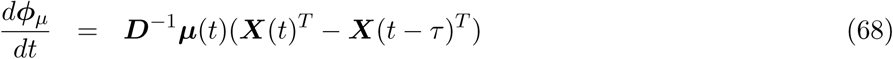

This learning rule is local in the space ***μ***(*t*) as matrix ***P*** (*τ*) is now replaced with the diagonal matrix *D^−^*^1^. Furthermore, the diagonal elements of this matrix are ***D**_ii_* = (‖***μ***_*i*_(*t*)‖^2^ +*λ*)^−1^ where ‖ ‖ is the standard *L*_2_ norm. This is what motivates us to refer to this learning rule as the Fourier rule, as this dynamic *L*_2_ optimization with an oscillatory ***r***(*t*) projects onto an orthogonal (but incomplete) basis ***μ***(*t*) of the 𝕃_2_ function space. This is the idea behind the generalized Fourier series in functional analysis. Now, the local Fourier rule is derived under the assumption that the spike train, ***r***(*t*) is very nearly orthogonal and the ***r***(*t*) have similar magnitudes in the *L*_2_ space:

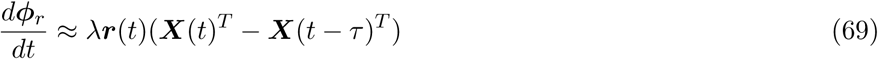

where *ϕ_ij_* only depends on ***X***_*j*_(*t*) and ***r***_*i*_(*t*) and *λ* is a constant. Finally, one can reduce the local Fourier rule to the following form:

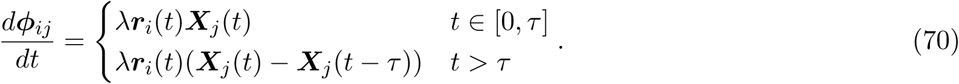

Replacing ***X***_*j*_(*t*) with a filtered post-synaptic spike train, *r_post_*(*t*), we obtain a modified version of the Hebbian learning rule:

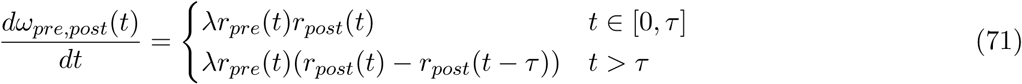

for the weight *ω_pre,post_* coupling a presynaptic neuron to a post-synaptic neuron. The key element in a Hebbian learning rule is the online computation of inner products. When combined with oscillatory spike trains, this allows for accurate online function approximation and the formation of memories in weights.

